# Dendrites support formation and reactivation of sequential memories through Hebbian plasticity

**DOI:** 10.1101/2023.09.26.559322

**Authors:** Alessio Quaresima, Hartmut Fitz, Renato Duarte, Peter Hagoort, Karl Magnus Petersson

## Abstract

Storage and retrieval of sequences require memory that is sensitive to the temporal order of features. For example, in human language, words that are stored in long-term memory are retrieved based on the order of phonemes. It is currently unknown whether Hebbian learning supports the formation of memories that are structured in time. We investigated whether word-like memories can emerge in a network of neurons with dendritic structures. Dendrites provide neuronal processing memory on the order of 100 ms and have been implicated in structured memory formation. We compared a network of neurons with dendrites and two networks of point neurons that have previously been shown to acquire stable long-term memories and process sequential information. The networks were equipped with voltage-based, spike-timing dependent plasticity (STDP) and were homeostatically balanced with inhibitory STDP. In the learning phase, networks were exposed to phoneme sequences and word labels, which led to the formation of overlapping cell assemblies. In the retrieval phase, networks only received phoneme sequences as input, and we measured the firing activity of the corresponding word populations. The dendritic network correctly reactivated the word populations with a success rate of 80%, including words composed of the same phonemes in a different order. The networks of point neurons reactivated only words that contained phonemes that were unique to these words and confused words with shared phonemes (success rate below 20%). These results suggest that the slow timescale and non-linearity of dendritic depolarization allowed neurons to establish connections between neural groups that were sensitive to serial order. Inhibitory STDP prevented the potentiation of connections between unrelated neural populations during learning. During retrieval, it maintained the dendrites hyperpolarized and limited the reactivation of incorrect cell assemblies. Thus, the addition of dendrites enables the encoding of temporal relations into associative memories.

## Introduction

Speech perception results from the integration of continuous streams of acoustic information over time. Understanding how this capacity is grounded in the underlying neural activity remains a challenge for the brain sciences. One aspect of the phenomenon to be explained concerns how the brain accesses long-term memories on the base of environmental stimuli with temporal extent. During spoken word recognition, it corresponds to recollecting words from a lifelong learned lexicon based on the phonological information (McQueen & Gaskell, 2007). Although many of the neural markers of this process are known, a mechanistic theory of how word memories are learned, maintained, and accessed is lacking (Poeppel & Idsardi, 2022). One hypothesis is that word memories are stored in the long term in the mental lexicon in the form of cell assemblies that are acquired through Hebbian learning (Pulvermuller, 1999; Garagnani *et al*., 2009); this view is in agreement with cognitive and computational theories of cortical processing based on the synaptic junctions (Amit, 1995; Fuster, 1997; Gastaldi *et al*., 2021). However, because of the *phonological overlap* between words in the human lexica (words being composed of the same phonemes, such as /kæt/ and /tæk/), accessing the word forms entails that cortical memories are sensitive to the order of the phonemes in the stimulus. According to long-standing critique of associative memories, these computational requirements cannot be achieved in the synaptic connections because associative memories lack the necessary expressiveness to encode relationships (Gallistel & King, 2011; Gallistel, 2021), i.e., the order among phonemes in human words. Thus, the main theoretical issue concerning the implementation of a biologically constrained word recognition model is the capacity of a spiking neural network to discriminate between sequences of similar inputs. We refer to this problem as the sequence detection problem. In the next three sections, we outline the requirements for long-term and short-term memory imposed by sequence recognition and finally present a model to solve this cognitive-computational problem.

### Formation and retrieval of word memories

A cell assembly is a functional population of neurons that emerges from the repeated co-activation of cells (Hebb, 1949). Reciprocal firing leads to synaptic potentiation and the formation of synaptic engrams in the form of interconnected subnetworks. The engrams are carved in the neuronal tissue through experience, following associative plasticity (Langille & Brown, 2018). Accordingly, assemblies are considered the core unit of memory (Poo *et al*., 2016). The strengthening of their synaptic connections, that is, the formation of an engram through long-term potentiation (LTP), is considered the causal landmark of memory formation (Dringenberg, 2020).

In the cell assembly view, the recognition of a familiar stimulus during sound perception corresponds to the reactivation of groups of auditory neurons and generates a burst of activity (Sakurai *et al*., 2018). Reliable, correlated cell activity is widely observed in the experimental literature on human and non-human animals (Almeida-Filho *et al*., 2014; Cohen *et al*., 2020; Hemberger *et al*., 2019), for a critical review of this phenomenon see Langille & Gallistel (2020). Compelling evidence on word recognition comes from two studies that obtained intracortical recordings in humans. Chan *et al*. (2014) observed bursts of neural activity in localized populations of the temporal cortex upon acoustic presentation of single words, but not when the stimulus was played backwards. Secondly, Vaz *et al*. (2020) isolated neuronal bursts in the middle temporal gyrus during recall of spoken word associations. Cortical neurons reactivated in ordered sequences during the learning period and when the cue-target pair was correctly recollected, but they lacked sequential structure when the association was wrong. Taken together, these studies suggest that accessing spoken word memories entails sequential firing of cell assemblies in speech areas.

Sequential spikes within assemblies, named phase-sequences, were already indicated as one of the distinctive features of memory access by Hebb (1949). Sequential firing is pivotal for the passage of information to downstream neural populations with causal consequences on behavior (Buzsáki, 2010). Several models have shown how Hebbian plasticity leads to the generation of phase-sequences in spiking neurons; networks can be trained to repeat sequences and activate assemblies in succession (Shouval, 2011; Scott & Frank, 2023; Maes *et al*., 2020; Cone & Shouval, 2021; Haga & Fukai, 2018). These models are based on two properties of cortical learning. First, memories can be instilled in the network connectivity via the repeated presentation of the stimuli (Litwin-Kumar & Doiron, 2014; Zenke *et al*., 2015; Tomasello *et al*., 2018). Second, Hebbian plasticity, in the form of spike-time-dependent plasticity (STDP), supports the formation of directed connections, and thus sequentially sensitive activity (Clopath *et al*., 2010; Fiete *et al*., 2010). However, the fact that engrams can be arranged to produce sequential activity does not explain how such spike sequences can be read out from downstream neural assemblies.

The problem becomes pressing if one intends to explain word recognition based on the neural mechanism of cell assembly; the order of phonemes in phonological overlapping words is key to distinguishing between word memories. In this respect, two questions must be addressed. First, can sequential activity trigger the recollection of order-dependent memories based solely on associative synapses? Second, does Hebbian associative synaptic plasticity support the acquisition of such memories? On the one hand, detecting sequences based on synaptic connections is achievable; networks with fine-tuned asymmetries in the weights structures can re-activate assemblies based on the order of the stimuli (Sequence Detector Network, Knoblauch & Pulvermüller, 2005) and supervised plasticity rules render neurons sensitive to the order of pre-synaptic firing (Tempotron neuron, Gütig & Sompolinsky, 2006). On the other hand, these solutions have not been demonstrated to be valid in realistic conditions. For example, when the STDP rule was tested in a sequence labeling task in a biologically constrained network, plasticity among excitatory cells offered only a moderate contribution (Duarte & Morrison, 2014). Thus, it remains unclear how to induce order-sensitive memories in recurrent spiking models using Hebbian-like plasticity.

### Integration of long-term and short-term phonological memories

One possibility is that biological networks require additional computational primitives to perform sequence detection. For example, variables to carry information forward on the time scale of the memory to be accessed (Chaudhuri & Fiete, 2016). Theoretical and experimental work indicates that short-term memory (STM) of phonological information must be held in the system by a dedicated mechanism, distinct from long-term memory (LTM) (Norris, 2017). The volatile auditory information storage must also allow for learning of the new association in the LTM. Arguably, with a compatible STM mechanism, the STDP rule should be able to encode time-structure sensitivity network’s engrams.

We hypothesize that the STM must satisfy certain requirements in order to achieve sequence labeling. First, it must encode time information, for example, in its slow decay to the resting state; second, there has to be a silent memory such that rapid compensatory mechanisms do not erase it (Zenke *et al*., 2015); and, third, it should interact with the long-term rule in the network to induce potentiation of the salient synapses. Of the three main STM proposals in the computational literature, persistent activity (Wang, 1999; Papoutsi *et al*., 2014; Wang, 2021), synaptic short-term memory (Mongillo *et al*., 2008), and neuronal memory (Fitz *et al*., 2020; Salaj *et al*., 2020); the constraints described appear to be satisfied only by synaptic short-term memories. Crucially, few computational studies have tested the interaction of this type of STM with the formation of cell assemblies. A previous study by Cone & Shouval (2021) demonstrated that networks can learn sequences on the timescale of human words using synaptic decay to encode STM and eligibility traces to update long-term memories. For each neuron, the synaptic memory was shared across pre-synaptic cells, which is not realistic considering the actual geometry of synaptic arrangements. In our opinion, the slow, post-synaptic decay that was implemented is better expressed by regenerative events in segregated dendritic compartments, for example, the NMDA receptor spikes. The crucial role played by dendritic processes in detecting spatiotemporal sequences is indeed demonstrated both experimentally and computationally (Branco *et al*., 2010; Bhalla, 2017).

We propose that dendritic memory (Quaresima *et al*., 2022) can satisfy these requirements as well as support the formation and maintenance of sequence memories on the timescale of human spoken words. Dendritic memory is expressed by long-lasting (100 ms) plateau potential, elicited by NMDA receptor spikes; the dendrites undergo long depolarized states that can bind the sequences of incoming phonemes. In addition, because the dendritic compartments are segregated, and the somatic firing activity is only weakly coupled to the dendritic membrane potential, allowing for the silent encoding of short-term memories. The hypothesis is that if STM is expressed as dendritic memory, it will induce the formation of long-term memories through vSTDP, thus supporting sequence detection in biological networks.

### The Tripod network model and the sequence recognition task

Dendritic neurons are modeled as Tripod neurons (Quaresima *et al*., 2022), a three-compartment neuron model with two dendritic compartments and a soma. The dendritic compartments are endowed with NMDA receptors and voltage-gated channels that allow for dendritic memory in the order of a hundred milliseconds (Fig.1*A*). The memory is provided by the long-lasting dendritic depolarization following an NMDA spike.

**Figure 1:**
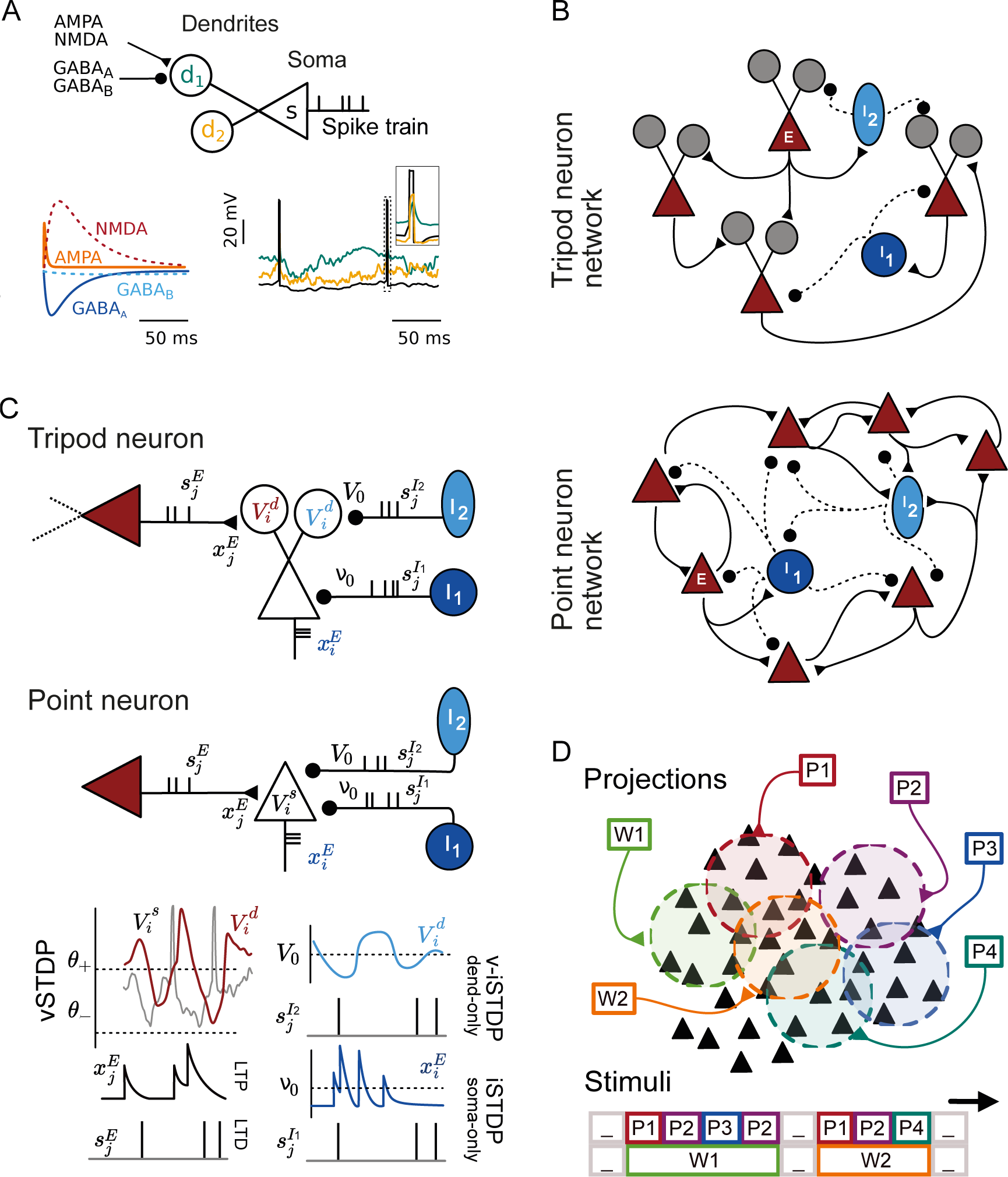
Tripod network: neuron models, connectivity, and plasticity rules. **(A)** The Tripod neuron has two dendritic compartments (circles) and a soma (triangle). Dendritic length determines the electrical properties of the compartment. Synapses included glutamatergic (AMPA, NMDA) and gabaergic (GABA_A_, GABA_B_) receptors with the illustrated timescales. The soma was modeled as an adaptive-exponential neuron and somatic spikes were backpropagates to the dendrites, as shown in the inset. **(B)** In the Tripod network, excitatory neurons targeted the dendrites of other excitatory neurons and the soma of inhibitory neurons. Inhibitory neurons I1 (fast-spiking) and I2 (adaptive) targeted the soma and dendrites of Tripod neurons. They also form recurrent connections within the inhibitory populations. In the networks of point neurons, all synapses connected to the somatic compartment. **(C)** Schematics of excitatory and inhibitory synapses plasticity. The left side illustrates voltage-based STDP in the glutamatergic synapses between excitatory neurons (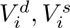 are the membrane potential of dendritic and somatic compartments). The plasticity rules are the same in the dendritic and somatic compartments, except for the thresholds of long-term potentiation (LTP) and depression (LTD), respectively *θ*^+^ and *θ^−^*. Right side; the inhibitory synapses onto the excitatory cells are subject to iSTDP. The equilibrium values depend on the neuron types. I2cells aim to stabilize the dendritic potential at *V*_0_ = −70 mV (v-iSTDP) and I1cells regulate the firing rate of the soma, *x^E^* = 10 Hz (iSTDP). Both excitatory and inhibitory plasticity is driven by the pre-synaptic activity, in the form of filtered spike train 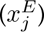 or sum of delta functions 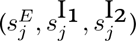 **(D)** External projections target a subset of excitatory neurons. The phoneme and word projections are fixed and target 5% of the excitatory population in both dendritic and point-neuron models. The phonemes are stimulated in sequence, and the corresponding word is stimulated throughout the interval.

The network is composed of 2000 excitatory Tripod neurons, 175 fast-spiking interneurons, and 325 slow-spiking inhibitory neurons, modeled as point neurons (I1, I2). Each neuron is connected to 20% of the other neurons in the network. The synaptic connections onto Tripod neurons are located on the dendrites (Fig.1*B*) and subject to voltage-dependent spike-timing-dependent plasticity (vSTDP, Clopath *et al*., 2010; Bono & Clopath, 2017). Excitatory plasticity supports the formation of engrams by strengthening the post-synaptic connections of neurons that fire onto depolarized dendrites. Because of the strongly connected cell assemblies, the network is prone to runaway activity. We used two additional mechanisms to prevent this from happening. First, homeostatic plasticity keeps the total incoming synaptic strength constant (multiplicative synaptic scaling, Tetzlaff *et al*., 2011). Second, the network is stabilized by means of fast compensatory mechanisms in the form of inhibitory spike-timing-dependent plasticity (iSTDP, Vogels *et al*., 2011). iSTDP is implemented via plastic connections between I1 neurons and the soma of the excitatory cells. In addition, we introduce a voltage-dependent iSTDP rule between the I2 population and the targeted dendritic compartment to reach a balanced excitatory-inhibitory synaptic input on the dendrites (Fig.1*C*).

We investigate word recognition in a Tripod network simulation. Word memories are intended as long-term associations between sequences of phonemes and words assemblies. The assemblies are induced on randomly selected subsets of the excitatory population, with word and phoneme assemblies loosely overlapping (Fig.1*D*). We use external input projections to induce the strengthening of synapses within and across the cell assemblies. The projections stimulate the dendritic compartments.

The network simulation is divided into two phases. The first phase is the associative one and we present both words and phonemes stimuli, simultaneously. On the base of the STDP rule among excitatory neurons, the co-activation of phonemes and word populations is expected to form auto-associative (recurrent engrams) and, possibly, hetero-associative (feedforward connections, between phonemes and words) memories. In the second phase, the recall phase, we turn off the plasticity mechanism and present only phoneme sequences. During recall, we measure the activity of all word populations and consider word recognition successful if the target word population is the most active one in the pool of the word populations.

We test word recognition on seven lexica that differ in the number of words they contain (8 to 17) and the amount of phonological overlap among the words. Two of them, *Identity* and *No overlap*, contain words whose phonemes are not shared across the words in the lexicon; thus, each phoneme is univocally associated with a word. Conversely, the other lexica comprise words with phonological overlap. An example from the lexicon *Overlap* are three words *log, dog,* and *god*; the first two words share two phonemes, but the second and the third share all the phonemes, and they are distinguishable only on the base of the order of the phonemes presented. Thus, correct word selection requires phonemes-word associations sensitive to the order of the phonemes in the stimulus rather than their sole identity features. The network must create order-sensitive associations to achieve correct word recognition. We compare the Tripod network with two point-neuron models endowed with similar plasticity mechanisms and measure the differences in the recognition score, its dynamics, and the synaptic structure formed following the associative phase.

## Results

We investigated the capacity of the Tripod network to form and recall time structure-dependent memories in a word recognition task. Because the dendritic neuron model has not been studied in a recurrent network before, we first set the network to a realistic operational point and verified the absence of pathological dynamics. We tuned background noise and synaptic strengths such that the network was in a regime of sparse firing. When the three synaptic learning rules apply (v-STDP, iSTDP, v-iSTDP), the network’s baseline activity has a low firing rate and tends to synchronize in slow bursts of activity. Conversely, when receiving external input, the network shows a low degree of synchrony and a sparse firing rate (Appendix A). The analysis indicates that the firing rate is heterogeneous across cells, both in the associative and recall phases: neurons receiving external projections fire more than neurons that do not. In the associative phase, a strong response from both phonemes and word populations is expected because of the external stimuli on both populations. Conversely, in the recall phase, word assemblies are not stimulated, and their activation originates from the reverberation of activity in the phoneme populations.

The following sections analyze in detail the activation of word assemblies in the recall phase. First, we show that for each sequence of phonemes, the most reactivated population is the one corresponding to the word associated with the phoneme sequence (the word *doll* and the sequence *D, O, L, L*). Then, by using the firing activity of word assemblies as the index of word recognition, we estimate the network capacity to detect sequences in a set of lexical with increasing phonological overlap. In the remaining sections, we investigate the mechanisms that allow the network to recall word memories. To determine the role of the dendrites in the network, we test four additional networks of point-neuron and dendritic models on the same task and evaluate their recognition capacity and dynamics. Thus, we analyze the network structure that emerges during the learning phase. We demonstrate that the phonemes-to-word connections strengthen (long-term storage) as well as that the dendritic non-linearity (short-term memory) determine the network’s capacity to recognize sequences.

### Word assemblies are sensitive to phonological order

We start the analysis of the network recognition capacity by taking a closer look at the coordinated firing of word and phonemes population during the associative and recall phases. To highlight the structure of the network activity, we ordered the network neurons by the projections they receive and plotted their spikes in the raster plots in Fig.2*A*. The figure shows the network activity for the eight phonemes and ten words in the lexicon *Overlap* during both phases. In the recall phase, only phonemes populations receive external inputs. The lexicon used in this experiment includes words with phonological overlap. Some words are contained within others (e.g., *poll*/*pollen*), some are anagrams of one another (e.g., *lop*/*poll*) or reversed (*dog*/*god*).

**Figure 2:**
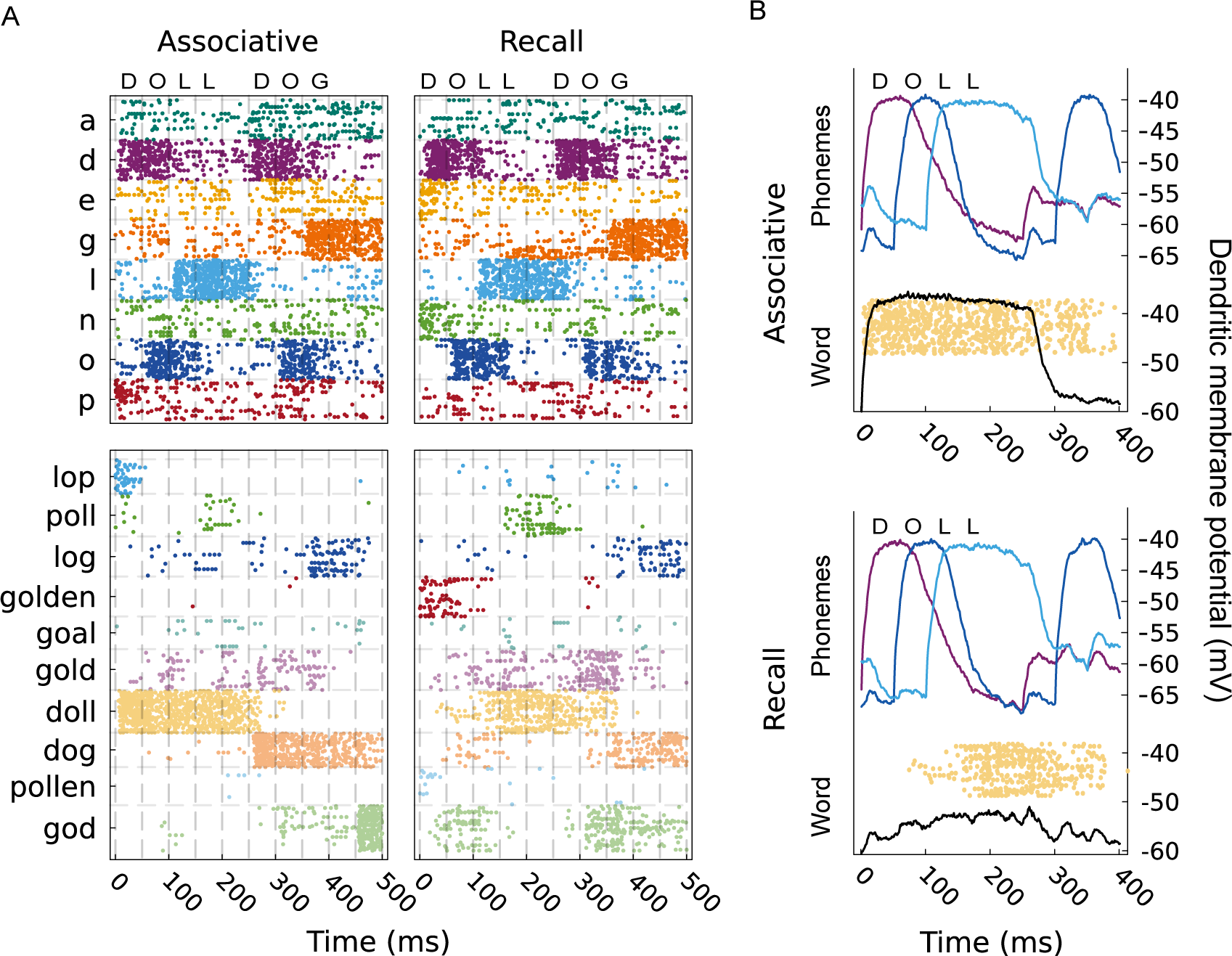
Network activity in the associative and recall phases. (**A**) Raster plot of phoneme and word populations during the presentation of the words *doll* and *dog*. Individual phonemes are stimulated for 50 ms each with a silent interval of 50 ms between words. In the associative phase, the word assemblies are stimulated for the entire interval in which the words are activated (*doll* 200 ms, *dog* 150 ms) (left). Conversely, they are not stimulated in the recall phase but activate from the reverberation of activity related to phoneme associated populations (right). (**B**) Average dendritic membrane potential for the assemblies associated with the phonemes *D, O* and *L,* and the word *doll*; the panels show the associative (top) and recall (bottom) phases. In the associative phase, the external projections strongly depolarize the phonemes and word assemblies. In contrast, in the recall phase, only the phonemes reach the −40 mV depolarization. The membrane potential of the word assembly slowy builds up and reach the maximum depolarization at word offset. Similarly, the firing activity of the word population changes in the two phase. In the associative phase the word assembly starts firing at word onset (the yellow dots indicate the assembly’s spikes), conversely, in the recall phase, the word population fires only after 50 ms to 100 ms from the onset of the phoneme sequence.

We first analyze the activity of the assemblies in the associative and recall phases. The raster plot shows that the activity of phoneme assemblies lasts longer than the stimulation intervals (grey vertical lines). Phonemes are stimulated for 50 ms only, but their firing persists for about 50 ms after the offset of the stimulus. Although the external input was the same for all phoneme populations (8 kHz for 50 ms), phoneme assemblies were not reactivated with the same strength. For example, the phonemes *D*, *G*, and *L* respond more strongly to the external inputs than the phoneme *O*. Comparing the activity across the entire simulation, it appears that the average activity of the phoneme assemblies depends on the total number of occurrences of the phoneme (App. Fig.1). The low firing of frequent phonemes is due to both neuronal adaptation, which tends to hyperpolarize the frequently stimulated populations, and the homeostatic component of the plasticity rule, which penalizes recurrent connections associated with frequent phonemes. Like phoneme assemblies, the word populations’ neurons fire in the two phases. In the associative phase, the word populations are activated along with the phonemes, and the external stimulus dominates their time course. The beginning of the assembly firing coincides with word onset and quickly fades when the following word is presented. In the associative recall phase, the word populations do not receive input but are nonetheless activated by the activation of phoneme populations; they receive the external stimulus through the reverberation of the phoneme assemblies. All words that contain the input phonemes in some position are partially reactivated. For example, for the input sequence *D, O, L, L*, the words *dog*, *poll*, and *gold* are activated, however the population of the word *doll* fires more; presumably because it matches both the identity and the order of the phonemes (Fig.2*A*). This behavior indicates that the word assemblies activate in a manner that depends on the order of inputs and that the network can detect sequences.

Before proceeding with the analysis of the sequence recognition capacity, we explored the dynamics of the dendritic membrane potential of phonemes and word assemblies. Fig.2*B* offers insights into the prolonged activity of the assemblies. The panels illustrate the membrane potential of three phonemes (D, O, L) and the word assembly *doll* during the presentation of the corresponding phoneme sequence (*D, O, L, L*). The dendritic membrane potentials of both the phoneme- and word populations remain depolarized beyond the duration of the stimulus. Such slow decay is supported by the dendritic memory of the Tripod neuron, and it is due to the rise of NMDA spikes in segregated dendrites. In the intervals in which the phoneme population activity overlaps with the depolarized word population, the synaptic connections are potentiated by the STDP rule. Despite the average word membrane being below the LTP threshold, certain cells are sufficiently depolarized and form stable connections within and between assemblies. Thus, the large dendritic depolarization due to stimulation enables the formation of phonemes-to-words associations that are sensitive to the co-activation of phonemes.

In the recall phase, the reactivation of the target word population takes place gradually during the presentation of the phonemes. Because the NMDA spikes are all or none events in which the membrane potential reaches approximately −20 mV, the slow building up of dendritic potential in the lower panel of Fig.2*B* indicates the recruitment of more neurons in the word’s assembly. The slow depolarization transforms into a burst of activity after 100 ms, when enough neurons are depolarized. The consequence of this dynamics is that the peak activity of the word population lags behind the onset of the phonological stimulus. Such delayed responses are observed across all the words inspected, and it appears to be an intrinsic property of word assembly reactivation in the dendritic model. In the following, we individuate an optimal interval in which word populations are maximally activated and use this interval to quantify the word recognition in the model.

### Word recognition latency

The previous analysis indicates that in the Tripod network word populations are reactivated by the activity in the phoneme assemblies. In addition, the response of the target word population has a latency of approximately 100 ms. We now define a measure of word recognition that allows us to identify correct word recognition and accounts for the delay in word reactivation.

To start with, and elucidate how the measure works, we analyze the firing activity of the assembly associated with the word *doll*. The word lasts four phonemes, which corresponds to 200 ms. To inspect if the network reactivates the correct memory, we compare the assembly firing rate of the correct word population with those of the other populations during the interval in which the input is presented. For each interval in which the sequence *D, O, L, L* is presented, the corresponding word population should activate. Thus we can identify each of these intervals as trials to test word recognition. In addition, because of the word’s population latency, we also test an interval shifted forwards of 100 ms. An illustration of the comparison among word population rates is presented in Fig.3*A*; the eight panels show the average firing rate of the target population *doll* against five competitor words (*dog, god, log, poll* and *goal*) and across the 110 trials where the sequence of phonemes (*D, O, L, L*) was presented. The activity of *doll* is shown on the vertical axis of each panel, and the horizontal axes show the activity of the word competitors. The black circles present each trial, and the red dots indicate when the average firing across the entire simulation is significant. The number annotated on each panel is the chance-corrected Cohen’s (*κ*) word recognition score, based solely on the firing rate (Methods).

**Figure 3:**
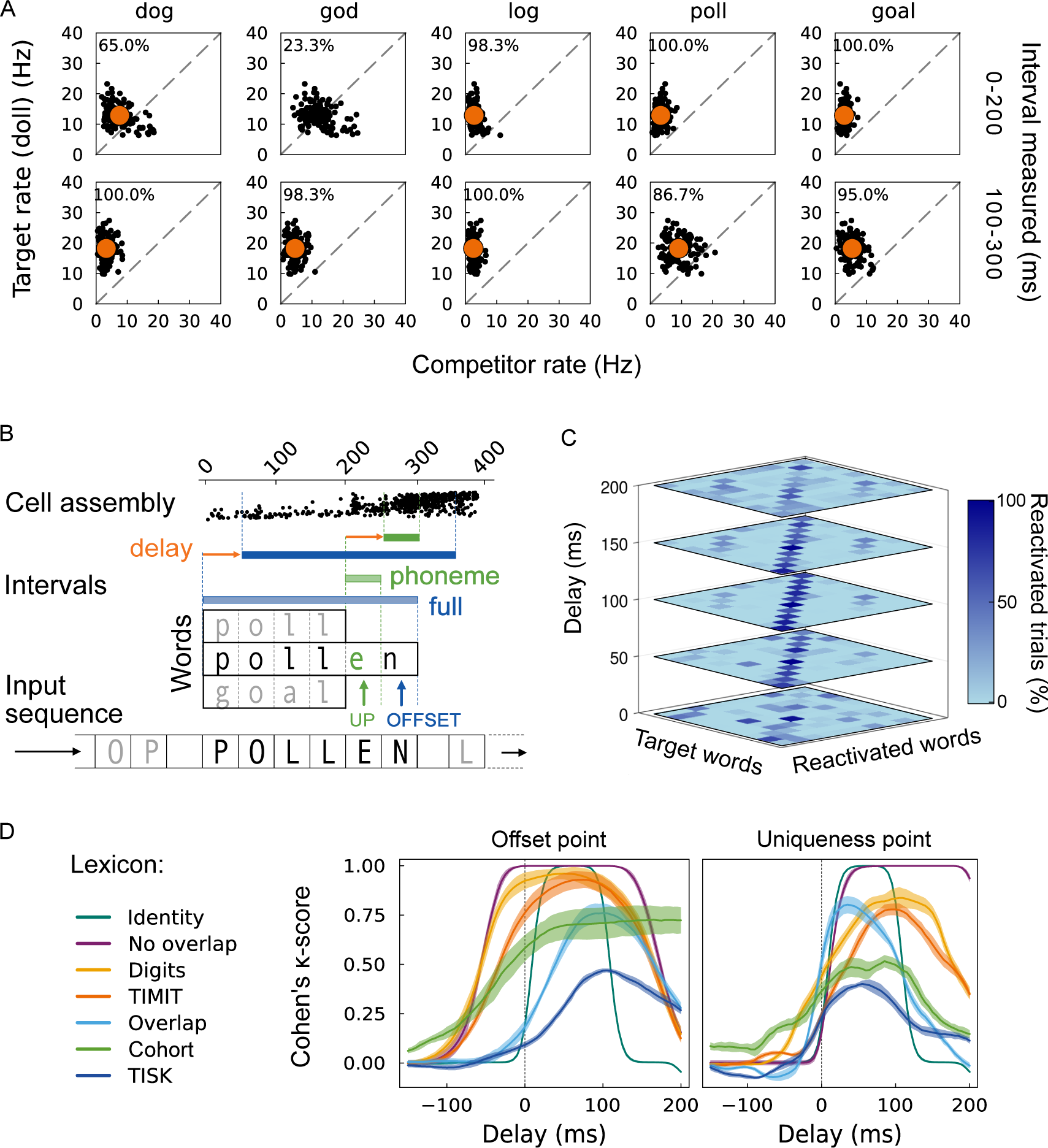
Recognition accuracy at word offset and uniqueness point. (**A**) Firing rates of the target word population against competitor populations for the phoneme sequence *P, O, L, L*. Rates are measured over the duration of the target word, shifted forward in time by 100 ms. Each dot represents one trial. When located above the diagonal, the target population was more active than the competitor word. Orange dots show the mean over 100 trials. (**B**) Word recognition is tested at the word offset and uniqueness points (blue and green arrows). Assembly activity is measured as the average firing rate in the interval between word onset and offset (full interval, blue bar) or the interval of the phoneme at the uniqueness point (single phoneme interval, green bar). The word with the highest firing rate in the measured interval is chosen as the network output. The measured interval is shifted in time (delay, orange arrow) to test the evolution of the network dynamics recognition time frame. (**C**) Network responses are summarized in the confusion matrix that indicates the percentage of trials in which a word was activated for each target word. Recognition accuracy is higher when the diagonal stands out, indicating that the correct word was reactivated for most of the trials. Off-diagonal values different from zero indicate failed recognition. The vertical axis shows the time shift at which the matrix was computed. The matrices that are shown were obtained for the *Overlap* lexicon at the offset point. (**D**) Cohen’s *κ*-scores for the seven lexica averaged across ten samples for the offset (left panel) and uniqueness (right panel) point measures, plotted as a function of the interval delay. The network reaches maximum accuracy for delays between 50 ms to 150 ms for all lexica. The sharp increase of the *κ*-score in the right panel highlights the fact that word recognition cannot be achieved before the uniqueness point.

When the assembly’s activity is measured in the same interval of the phonological input (0 ms to 200 ms, upper panel), the firing rate of the target is systematically less than when it is measured with a delay of 100 ms, in some cases, it is not even sufficient to distinguish between the target and the competitor (*god*, second panel). By taking the average on all words, including the remaining five, we estimate that when the word recognition measure is delayed 100 ms, the word assembly *doll* is correctly recognized in 90% of the trials, while only in the 60% when it is measured in the same interval of the phonological input. Because the delay is computed starting from the offset point, we refer to this as the offset point (OP) measure.

WHAT IS THE OP MEASURE? We can generalize the procedure described for the word *doll* and determine the optimal word recognition latency in the network for all the words. One issue is the difference in length among the lexical items; some words contain more phonemes than others. For example, the OP measure of *doll* computes the average firing over four phonemes, (200 ms), but its competitors have three, four, or six phonemes. This difference can introduce biases when comparing the firing rates of the associated assemblies. For this reason, we integrate the OP measure with a second one with a fixed length. We measure the firing rate for the duration of one phoneme (50 ms) and do so at the uniqueness point (UP) of the word. The UP is the phoneme that makes the word unique in the lexicon; it depends on the presence of other words with shared onset phonemes. As before, we shifted the interval forward or backward in time to test the timing of word reactivation. We refer to this shift as the interval delay. The UP and OP measures are illustrated in Fig.3*B* and described in further detail in the Methods.

For each trial, we select the word associated with the population with the highest firing rate as the retrieved lexical item. Then we use it to compute a confusion matrix. The confusion matrix indicates which word was expected (target) and which was retrieved (reactivated). Correct word recognition implies that the target and reactivated words are the same and that the matrix has most of its trials on the diagonal. An example of the evolution of the confusion matrix with the delay is portrayed in Fig.3*C*. The stacked heatmaps show the confusion matrix for the lexicon *Overlap* based on the uniqueness point measure. The bottom matrix corresponds to 0 ms of delay, and the pairing between target and reactivated words appears random. The reactivated words match the target more and more as the delay increases, until a maximum of 100 ms and start decreasing. This is somewhat unexpected because all words are optimally recognized with the same delay despite the lexicon containing words with 3–6 phonemes. Moreover, it is important to notice that the 100 ms interval overlaps with the presentation of the following word. From the confusion matrix, we computed the *κ*-score as described in the Methods, the *κ*-score for the *Overlap* lexicon is 0.77%

To characterize the model’s capacity to recognize sequences in the presence of shared phonemes, we compared the recognition accuracy on seven different lexica that varied in size, number of shared phonemes (phonological overlap), and average word length. Some of the seven lexica have been used in previous studies on word recognition. The lexica and their statistical differences are described in the Methods. For the first two lexica (Identity and No overlap), words can be identified based on the first phoneme because it is unique to each word. Thus, recognition does not require sequence memory, and we refer to these lexica as memoryless. The other five lexica have phonological overlap and require temporal integration of symbols in a sequence. We computed the *κ*-score on the seven lexica for the offset and uniqueness point measures and varying the delay between −100 ms to 200 ms. The accuracy measure associates every delay shift with an average recognition score. Thus, it quantifies the latency of word recognition. The recognition *κ*-score is plotted against the delay in Fig.3*D*.

In agreement with the confusion matrices, the *κ*-score increased when shifting the measured interval forward in time. For both the OP and UP measures (left and right panels), recognition accuracy peaked for delays in the range 50 ms to 150 ms. The offset point measure has a smoother ramp than the UP because it averages the firing rate on a longer interval. Recognition accuracy was generally higher for the OP measure than the UP measure. At the OP, the highest accuracy is above 70 % for all but the *TISK* lexicon. At the UP, the recognition is proportional to the full interval measure (OP) but slightly lower (the Pearson correlation between the two measures is 0.9). The similarity between the two measures indicates the network activity before and after the short UP interval contributes marginally to word recognition. Most importantly, the UP measure (right panel) shows that recognition is not possible ahead of the word’s uniqueness point; for delays below 0 ms, most lexica had a recognition score close to zero. The UP measure indicates that before the UP, the network was in a state of co-activation of multiple assemblies with insufficient information to discern between words (flat confusion matrix, *κ* ≈0). Accuracy increases towards the UP for lexica with longer words because some of these words can be ruled out already before the UP, resulting in a higher chance of recognition (non-flat confusion matrix, *κ >*0). For example, in the *Cohort* lexicon, the sequence *C, A, P, I, T* will exclude the words *capias, capillary* and *capistrate* although *T* occurs before the uniqueness point. We tested ten randomized realizations of the network, which reached similar accuracy on each lexicon. The standard deviation of the *κ*-score is below 5 % for all lexica and conditions, except *Cohort* in the OP measure where it reached 10 %.

### Dendrites are needed to recognize words with phonological overlap

To understand the computational role of dendrites in the formation and reactivation of word assemblies, we compared the Tripod network with four other models, two with dendrites and two without. The networks of point neurons were chosen from the literature for their biological plausibility and relevance to the task at hand. The dendritic networks are instead copies of the original model but stripped of the asymmetry in the dendritic compartments. By comparing the models, we aim to isolate the network mechanisms that support word recognition. We hypothesize that the dendritic memory, provided by electrical segregation and NMDA receptors in the dendritic compartments (Quaresima *et al*., 2022) is the computational primitive that supports the recognition of words with phonological overlap.

The first point-neuron model was the network described in Litwin-Kumar & Doiron (2014) (LKD for short). It has previously been shown to learn and maintain stable cell assemblies in the presence of ongoing plasticity and background noise, using interacting forms of excitatory and inhibitory STDP. Assemblies developed for 20 stimuli randomly injected into the network during the early phase, similar to our associative phase. Later, in the absence of inputs, the network spontaneously and robustly reactivated the assemblies, demonstrating that the network learned the memories presented in the early phase. The second model is based on the work of Duarte & Morrison (2019) (DM for short), which modeled an L2/3 circuit of mouse cortex. The original study investigated the role of various sources of heterogeneity on the network’s computational capabilities, such as temporal integration and delayed decision-making. The network had no plasticity in the synaptic connections, and its recurrent weights were calibrated to achieve a balanced excitatory-inhibitory state. Because word recognition requires associations between phonemes and words, we endowed the DM network with STDP on the synapses connecting excitatory neurons. Analysis showed that the DM network was unstable and prone to oscillatory dynamics. To stabilize the network, we added iSTDP and the v-iSTDP homeostatic inhibition, similar to the Tripod network but with both inhibitory plasticity mechanisms related to the soma of the excitatory neurons. Both the LKD and the DM networks implemented conductance-based synapses; only the DM network modeled NMDA receptors, which, independently from the results on dendrites, are known to play a role in working memory in that they support the onset of persistent activity in network cliques (Wang, 1999; Papoutsi *et al*., 2014; Wang, 2021). In addition, the two models differ in the density of their recurrent connectivity, the number of cells considered, and, most importantly, the presence of distinct classes of inhibitory neurons. The LKD model has 4000 excitatory cells and 1000 fast inhibitory neurons. Conversely, the DM has 2000 excitatory neurons, 175 fast-spiking gabaergic cells (I1) and 325 slow-spiking ones (I2). An illustration of the point-neuron models and their synaptic learning rules is shown in Fig.1*B* and *C*. Equations and parameters of the models can be found in the Methods.

Moreover, we compared the Tripod network with two other dendritic models, one with only one dendritic compartment and one model where the input and recurrent connections targeted both dendrites (symmetric network). The three models differ in their dendritic and afferent configurations but are all endowed with the dendritic memory provided by segregated compartments and NMDARs. Compared to networks of point neurons, the dendritic networks were more robust to parameter variations. Network stability is due to the location of the recurrent connections on the dendrites; because the axial conductance limits the maximum current flowing from the dendrites to the soma the networks are less prone to epileptic firing. The dendritic network configurations of the three models are illustrated in Fig.4*A* and further described in the Methods. Note that symmetry here refers to both the synaptic connectivity pattern and the dendritic lengths.

**Figure 4:**
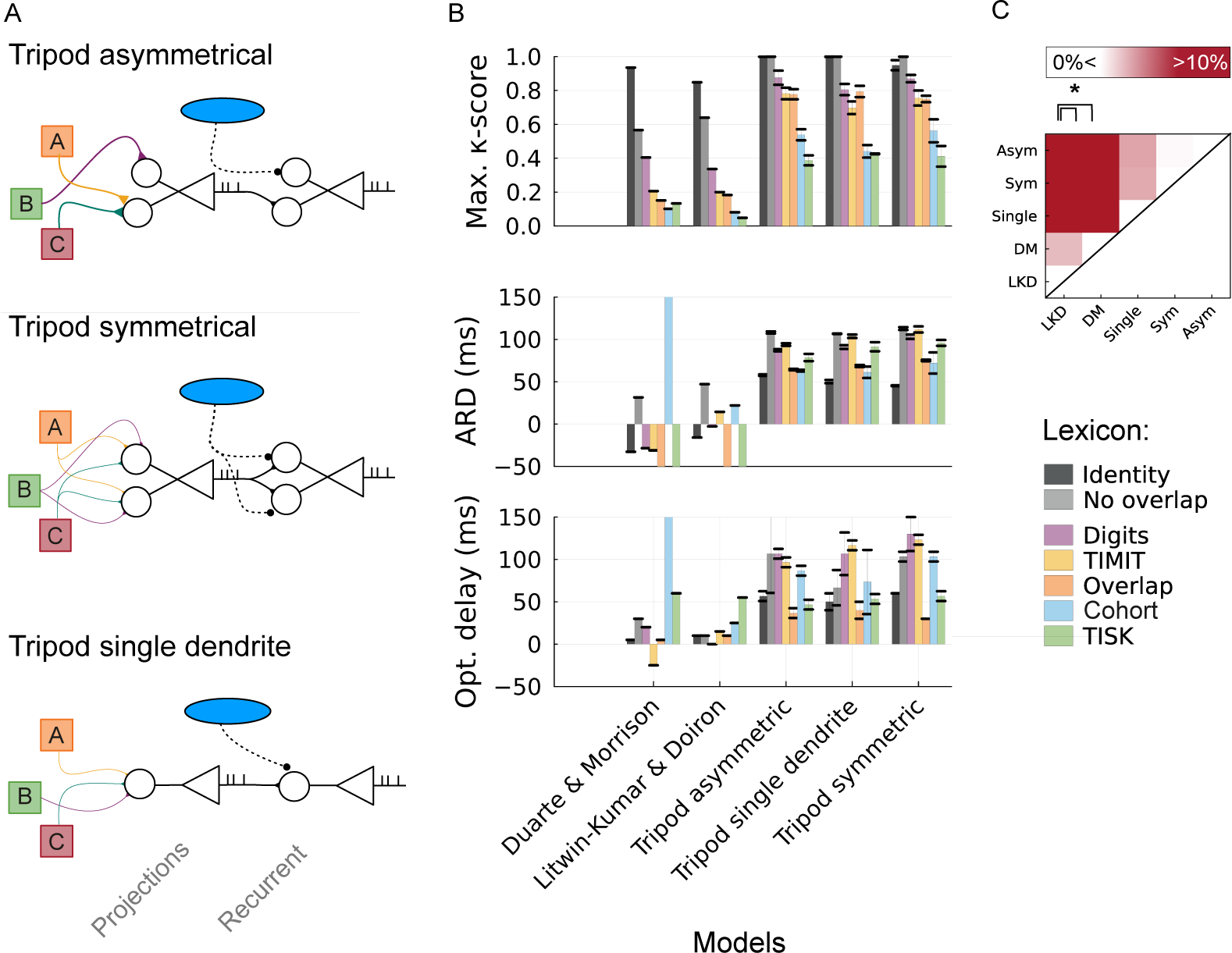
Comparison of point-neuron and dendritic networks. (**A**) Dendritic configurations and connectivity patterns of the Tripod-based network models. All three variants have dendrites whose length is uniformly random in the range of 150 µm to 400 µm. The asymmetric neuron has randomized pre-synaptic connections with density *ρ* =0.2 that target only one of the two post-synaptic compartments; the length of the dendrites is also drawn independently from each other. The symmetric network receives the same pre-synaptic connections, external and recurrent, on both dendrites, with half the connection strength than the asymmetric case; synapses have the same density (*ρ*); the two dendrites have the same length in the symmetric neuron model. Finally, the single dendrite neuron has only one segregated compartment, which receives all synaptic connections. **(B)** The three panels illustrate the maximal *κ*-score - across all delays, the average recognition delay (ARD), and their optimal delay (delay with maximum score) for the five network models and the seven lexica tested. Grayscale bars show the two memoryless lexica and colored bars show the five lexica that require temporal integration of phonemes for recognition. point-neuron models perform best on memoryless lexica with negative delays; it indicates that recognition is achieved before the offset point. Dendritic models outperform them on lexica with overlapping phonemes and reach high accuracy with a delay between 50 ms to 100 ms from the uniqueness point. The dendritic models’ average and standard deviations refer to a sample of three independent initializations per model. (**C**) The matrix shows the difference in accuracy between a model on the y-axis and a model on the x-axis, with a resolution of five percentage points, for the five lexica that require memory for word recognition.

The five models were exposed to the seven lexica introduced in the previous section. For each pair of models and lexicon, we computed the recognition accuracy according to the OP and UP measures, with delays in the range of −50 ms to 150 ms. We introduce two novel measures to elucidate the time course of word recognition: the average recognition delay (ARD) and the optimal delay. The ARD is a linearly weighted average of the delay intervals, using the *κ*-score as weight (Eq.18, Methods); the optimal delay is instead the delay with which the model reached the maximum score on the lexicon. The comparison between the models’ accuracy, optimal, and average recognition delays in the word recognition task is shown in Fig.4*B*. The panels refer to the UP measure. The analysis for the OP measure is qualitatively similar (SI Fig.4). The greyscale bars show the two lexica that do not require recognition memory, while colored bars refer to lexica that do. The *κ*-scores for the latter group were averaged for each model and compared pairwise between models in Fig.4*C*. The red color intensity codes for differences within +10 percentage points between models on the y-axis and models on the x-axis, with the significance level of the relative differences between pairs of models indicated.

The top panel shows that dendritic networks have systematically higher recognition scores than point-neuron models in lexica with phonological overlap. The point-neuron models perform well on the memoryless lexica, but their accuracy drops when temporal integration is required. The confusion matrices (SI Fig.4 *A*) showed that, in the latter case, the networks reactivated a subset of the word assemblies for any input sequence, suggesting that the functional association between the phonemes and word populations was not established. The point-neuron models did not learn the sequential structure of word memories. In contrast, the three dendritic models recognized all the words, in all lexica; the errors were distributed equally among the remaining words (SI Fig.4 *B*).

In addition, the ARD and optimal delay measures reveal that the time course of recognition differs in the two models. Dendritic models have high recognition, on average, for delays in the range 50 ms to 100 ms from the offset point (middle panel). Conversely, the delays corresponding to the maximum score (bottom panel) span a large range and depend on the lexicon property. The ARD of the point-neuron models is negative and its optimal delay is negative or close to zero, which indicates that the correct word populations are maximally re-activated before the offset point. For the lexicon that requires memory, the delay measure of point-neuron models is not informative because their scores are close to the chance level. The analysis indicates that immediate access to word memories is successful only if the identity of the early phonemes is sufficient to disambiguate the word and access the correct word memory, as it happens for the first two lexica. Otherwise, if the early phonemes contain insufficient information, the recognition will be, at best, based on the marginal distribution of the phoneme’s identity over the lexicon. The probability that the first phoneme belongs to the correct word decreases with the increase in phonological overlap. In dendritic networks, the reactivation of word memories relies on the temporal integration of the inputs, confirmed by the large values (50 ms to 150 ms) of average and optimal delays. Phoneme activity is carried forward in the dendrite’s membrane potential and the word population fully activates when the last disambiguation piece of the input sequence is presented at the uniqueness point.

Concerning the role of asymmetry in the network, the present results indicate that its contribution is marginal. Averaged over three independent samples, the asymmetric Tripod model scored higher than the other two dendritic models, but the differences are less than 5 to 10% and not significant Fig.4*C*. The delay measures of the three dendritic models resemble each other in offset and uniqueness point measures. The models’ similarities indicate that dendritic memory carries information over time independently of the connectivity and dendritic configurations tested (Fig.4*A*). In addition, the difference in accuracy among the three models does not depend on the lexicon; they all decrease their recognition accuracy for the lexicon with larger phonological overlap, suggesting neither asymmetry nor the number of dendrites provides additional mechanisms to cope with the increasing phonological overlap. Finally, the lack of significant differences between the two point-neuron models indicates the presence of two classes of inhibitory neurons with distinct inhibitory plasticity rules (I1, I2) is not the core mechanism for the word recognition task at hand.

From the comparison between the five models, we deduce that the dendritic memory endowed by the NMDARs in segregated compartments is the mechanism that allows the network to solve the word recognition task. However, because sequence detection requires the interaction of shortterm and long-term memories, it is not yet clear if the point-neuron models fail to establish the hetero-associative connections or, rather, cannot perform temporal integration. In the following section, we analyze the network connectivity emerging from the associative phase in the five models and show that only networks with dendrites form strong associations between phonemes and word assemblies.

### Dendritic networks have strong connections from phonemes to words

The present section investigates the network structure that supports sequence detection. To perform the network analysis, we reduce the model’s connectivity to an effective matrix that accounts only for words and phonemes assemblies. We define the effective matrix (C) as the average connectivity between the cells belonging to two assemblies (Methods). The effective matrix is a non-symmetric square matrix computed from the learned connectivity matrix at the end of the associative phase. The C is composed of four blocks, the connections between word assemblies (C*^W^ ^→W^*), between phoneme assemblies (C*^P^ ^→P^*), from phonemes to words (C*^P^ ^→W^*), and from words to phonemes (C*^W^ ^→P^*). An example of the C matrix from the asymmetric Tripod model for the *Overlap* lexicon is shown in Fig.5*A*. The top-left quadrant shows the feedback connections from word assemblies to phoneme assemblies, and the bottom-right quadrant shows the feedforward connections from phonemes to words. The top-right and bottom-left quadrants show the connectivity within phoneme and word assemblies, with strong recurrent connections on the diagonal. For completeness, the effective connectivity matrix for the remaining models is shown in SI Fig.5.

**Figure 5:**
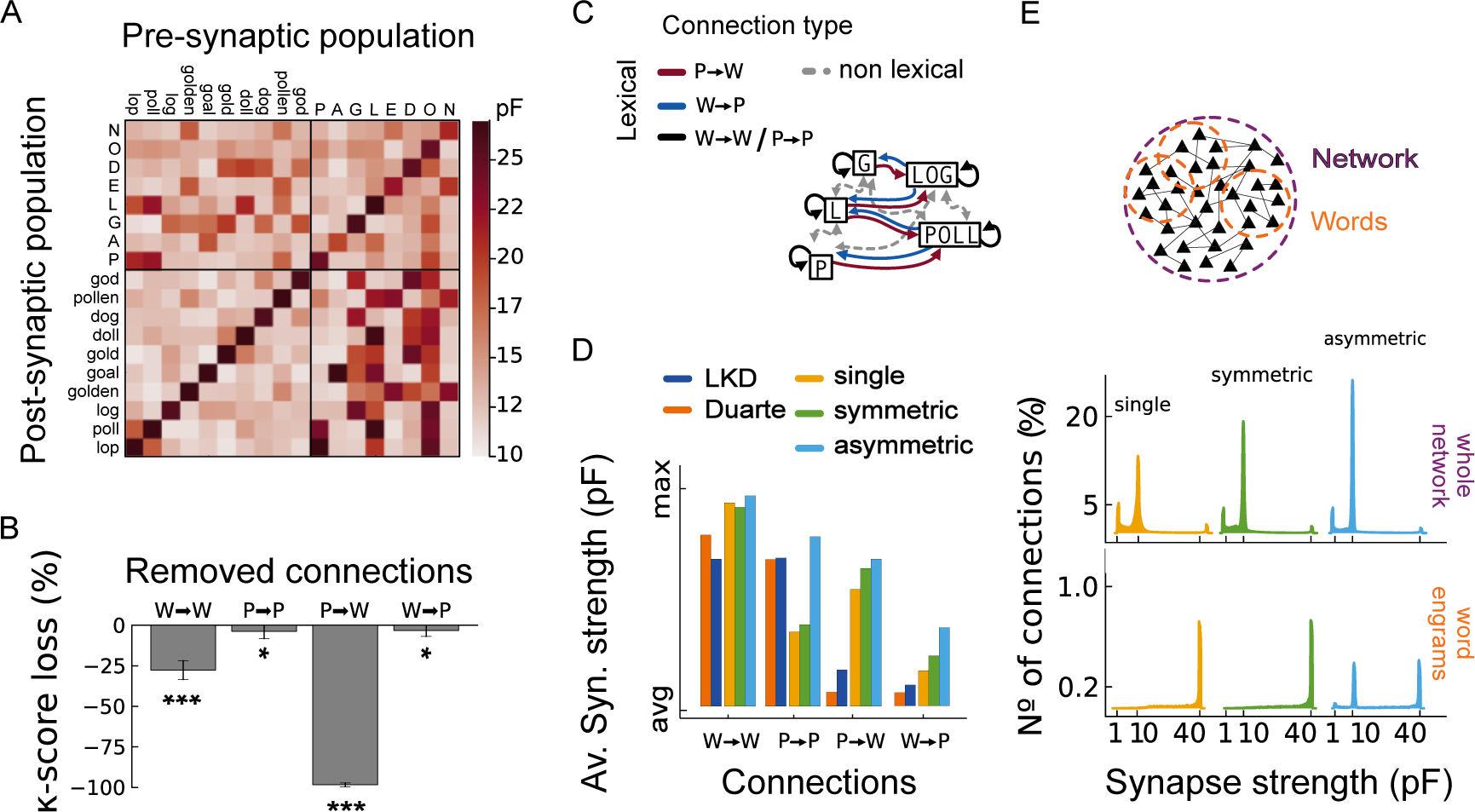
Feedforward structure in dendritic network support word recognition. (**A**) Effective connectivity matrix of the asymmetric dendritic network for the *Overlap* lexicon. The heatmap shows the average synaptic strength between assemblies, measured in pF. The four quadrants show the recurrent connections within phoneme and word populations, the feedforward connections from phonemes to words, and the feedback connections from words to phonemes. (**B**) The word recognition *κ*-score changes when each of the four quadrants of the effective connectivity matrix is leveled to the initial synaptic weight (the resulting matrices are shown in SI Fig.7). The four bars indicate the loss in recognition in each condition. They are all significant. However, removing the phonemes to word connection causes a drop in recognition of the 100%. (**C**) The schematic at the top illustrates the four types of lexical connections and separates them from the non-lexical ones, which are not expected to contribute to word recognition. The non-lexical connections bind phonemes with words that do not contain them or those among different words. (**D**) Average synaptic strength of the four types of lexical connections for the point-neuron and dendritic models. All models have strong connections within word assemblies, but only dendritic models have strong phoneme-to-word connections. (**E**) Distribution of synaptic strength over the entire network (top) and the word recurrent connections (bottom). The top panel shows that most synapses remain close to their initial strength of 10 pF, less than 3% of the connections developed stronger synapses (40 pF). The bottom panel (zoom in the range 0 to 1% of synaptic connections) indicates that for the symmetric and single dendritic models, the word assemblies have only strong recurrent connections. In contrast, in the asymmetric network, recurrent word engrams have both weak and strong synapses within the engram.

The four connection types of the effective matrix contribute differently to the word recognition task. To test their impact, we selectively removed each of the four blocks from C. The resulting effective matrices are similar to the original Fig.5*A*, except for the removed block, whose elements are set to the average connection strength (SI Fig.7). Thus, we measured the recognition score and calculated the percentage of recognition loss in each of the four conditions. The recognition loss is computed as 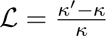, where *κ^′^* is the score with the modified matrix and *κ* is the baseline score. The loss is computed for each of the seven lexica and then averaged. The results of the ablation experiments (Fig.5*B*) indicate that the phonemes to word connections (C*^P^ ^→W^*) are pivotal for word recognition, recognition drops of the 100% when they are removed. Such connections are necessary because they relay externally-driven activity in the phoneme populations to the word assemblies. They emerge in the associative phase when phoneme and word populations are simultaneously activated. Interestingly, the phoneme-to-word connections for the *Overlap* lexicon have a similar, although weaker, structure in the DM network but are absent in the LKD network (SI Fig.5). The word-to-word connections are also important but not strictly necessary; in this case, the recognition drops of 25%. The other two blocks in Fig.5*A* involve synapses targeting phoneme populations. Their contribution to word recognition is marginal compared to the others, although the loss is significantly larger than zero. These connections have a smaller impact on the network dynamics because they are weaker than the external projections that stimulate the phonemes populations.

**Figure 6:**
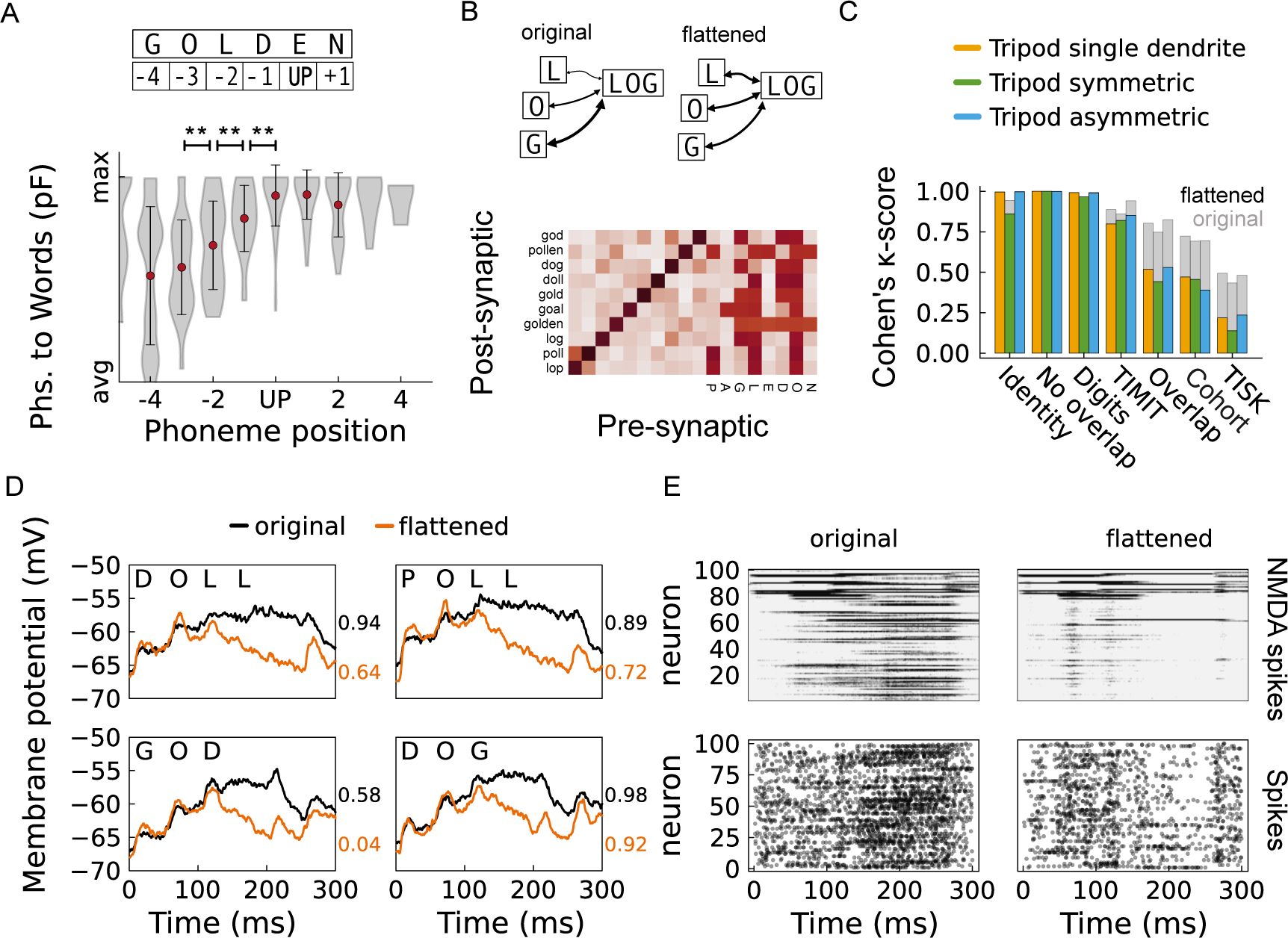
Feedforward structure in dendritic network support word recognition. (**A**) Strength of the phoneme-to-word connections organized by the serial order of phonemes in the word. The phoneme position is computed relative to the uniqueness point (top panel). The graph shows data pooled from all words in the five lexica that require memory. Red dots indicate averages, while the violin plots account for the entire sample. Significative differences in the average synaptic weights are evidenced between the three phonemes preceding the uniqueness point. (**B**) Lower quadrants of the connectivity matrix with flattened structure in the phoneme-to-word connections. The matrix shows the modified phonemes-to-word connections, the original being in panel Fig.5A. Flattened connections maintain the associations between phonemes and words but remove the internal structure necessary for distinguishing phonologically overlapping words. (**C**) Word recognition accuracy in networks with flattened connections, compared with the original connectivity matrix (grey). The *κ*-score decreases for the dendritic models when lexica have a large phonological overlap. (**D**) Average dendritic membrane potential with the original and the flattened connectivity. The membrane potential is portrayed for four distinct word assemblies. The black line indicates the normal dendritic dynamics, and the orange is the one in the flattened condition. The four plots show that in the flattened condition, the cell assemblies have a weaker sustained depolarization after word offset. (**E**) Assembly dynamics for the word *doll*, also comparing the original and the flattened network. The upper panels show the dendritic membrane potential, with black traits indicating potential above 20 mV (NMDA spikes). The lower panels show the somatic firing activity. Both the membrane potential and the firing activity are pooled over 10 samples of the sequence *D, O, L, L*. The panels show that the lack of sustained depolarization is due to fewer NMDA spikes and results in weaker somatic activity.

**Figure 7:**
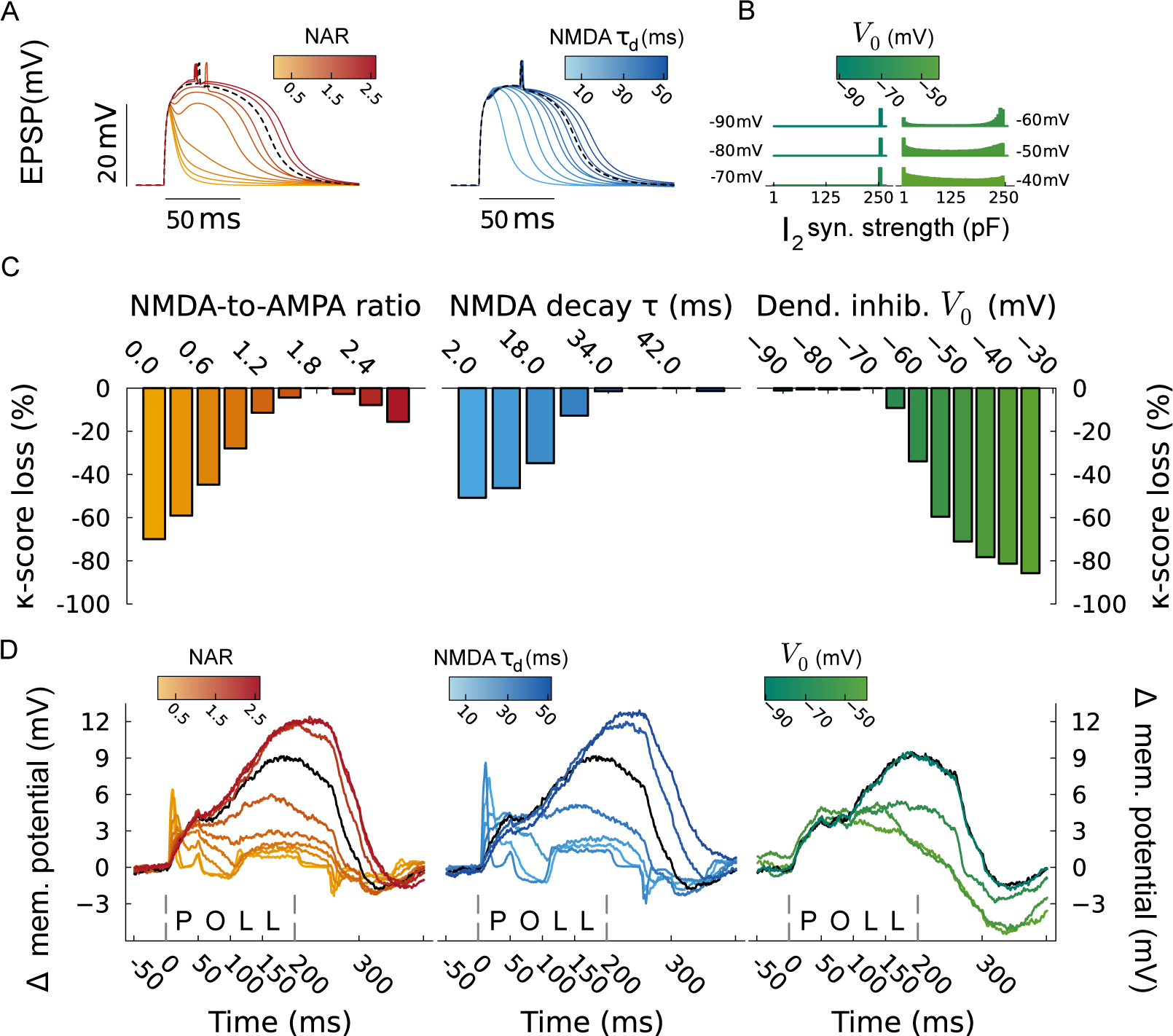
Dendritic non-linearity and inhibitory control enable word recognition. (**A**) EPSP of the dendritic membrane in Tripod neurons after stimulation of a single synapse. Curves show the membrane potential for varying NARs and NMDA decay timescales. The dashed line indicates the model that was used in the previous section, corresponding to a Tripod neuron with human synapses, with NAR = 1.8 and *τ_d_* = 35 ms. (**B**) Histograms of the inhibitory synaptic strength onto dendrites, measured at the end of the recall phase, for increasing v-iSTDP target potentials *V*_0_. Synapses potentiate towards their maximum value for *V*_0_ lower than −70 mV. (**C**) Loss of *κ*-score for variations in the parameters NAR, *τ_d_*, and *V*_0_. The bars with the darker shade correspond to the original model. Accuracy reduces up to 60 % when the dendritic non-linearity is absent (NAR < 0.6) or short (*τ_d_* < 18 ms). The changes are larger for increased values of the dendritic inhibition target potential (*V*_0_). (**D**) Difference in membrane potentials upon presenting the phonemes *P, O, L, L*. The plots express the difference between the average dendritic potential of the assembly associated with the word *poll* and the dendritic potential of the entire network. The difference in membrane potential is measured for all the parameter variations, i.e., NAR, *τ_d_*, *V*_0_. The black solid line is the same for the three panels and refers to the baseline asymmetric Tripod model.

Based on the outcome of the previous experiment, we inspected the effectivity connectivity matrix of the five models (point-neuron and dendritic network) to determine if the recognition capacity of the models was due to differences in C. To this aim, we calculated which phoneme and word populations should be associated for word recognition to succeed. We thus define a group of *lexical* connections as the connections from any phoneme to the words that contain it and vice versa, plus all recurrent connections within phoneme and word assemblies. The three groups that compose the lexical connections are illustrated in Fig.5*C*. We measured the average strength of the four types of lexical connections for the Overlap lexicon for the five models tested. To facilitate model comparison, the columns were scaled between the maximum average synaptic connection of each model’s C matrix (e.g., 26 pF in Fig.5*A*). The minimum corresponds to the initial synaptic strength of the excitatory to excitatory connections, which determines the synaptic weight budget for synaptic normalization. The initial weights are 10 pF for the dendritic models and vary for the two point-neuron models (DM: 0.45 pF, LKD: 2.76 pF). It was not possible to set the weights of the somatic models to be the same as the dendritic models because of instabilities in the network activity. We return to this issue in the Discussion.

Differences and similarities between the five models for the four lexical connection types are shown in the bottom panel of Fig.5*D*. For the dendritic models, the hetero-associative connections (phoneme-to-word) have comparable strength to auto-associative ones (words-to-words and phonemes-to-phonemes). In contrast, the point-neuron models have strong auto-associative synapses but weak connections binding phonemes to word assemblies. Interestingly, the recurrent phonemic connections are as strong as the word-to-word in the point-neuron and asymmetric models, but they are weaker for the symmetric and single dendrite models. All models have relatively weak word-to-phoneme connections. Overall, the analysis of the effective connectivity matrices indicates that the presence of dendrites promotes the strengthening of non-recurrent connections between phoneme and word assemblies. An overview of the effective connectivity matrices for the five models on all the lexica is presented in SI Fig.6. Comparing the panels across different lexica indicates that the formation of hetero-associative connections is achievable in point-neuron models if the words have no phonological overlap. A comparative analysis of these four types of connections is also presented in Appendix B, which analyzes the learning dynamics of the five models during the associative phase. the results are fully consistent with the ablation study presented here.

Further differences between the dendritic models can be gleaned from an analysis of the synaptic weights landscape of the three networks. The histograms in Fig.5*E* show the percentage of connections (y-axis) against their respective synaptic strengths (x-axis) for the entire network (top) and within the recurrent word engrams (bottom, grey shades show the entire network for comparison). The top row indicates that the dendritic models reach similar network configurations through Hebbian learning. The distributions of synaptic strength each have three peaks at 1.78, 10, and 40 pF. The two extrema are the minimum and maximum synaptic strengths allowed in the plasticity rule (Methods), and the mid-value is the synaptic strength with which the model is initialized. Synaptic normalization (i.e., homeostasis) enforces that if a compartment develops a strong incoming synapse, some other synapses will get weaker; one fully potentiated synapse requires four fully depressed ones. Thus, homeostasis explains the leftmost peak as a consequence of the strengthening of lexical connections. The most pronounced peak occurs at 10 pF. These synapses remain unchanged by learning and do not contribute to any engram. The asymmetric Tripod network has roughly twice as many idle connections as the single dendrite model and 40 % more than the symmetric one. The difference is visible in the lower panel; for the single and symmetric Tripod models, most of the recurrent word connections are fully potentiated, whereas only half are for the asymmetric model. Comparing the panels in Fig.5*D* and *E* with the effective connectivity of the symmetric and single dendrite network SI Fig.4, we infer that the asymmetric configuration recruits fewer synaptic resources for the recurrent word-to-word connections. The STDP redistributes the synaptic budget among the other connection types. The larger availability of synaptic resources explains why the asymmetric model has phoneme-to-phoneme and word-to-phoneme synapses whose strength is comparable to the word-to-word connections, which is not the case in the other two dendritic models.

### Structures in the effective connectivity matrix mediate recognition of words with phonological overlap

Because the phoneme stimuli do not encode temporal information, i.e., the position of the phoneme in the word, the network must rely on recurrent connections to process sequential order and reactivate the correct word assemblies. For example, the words *dog* and *god* share the same phonemes (lexicon *Overlap*), but their order is different. When presented with the phoneme sequence *D,O,G*, the word assemblies *dog* and *god* should assume different states such that sequential order information triggers the reactivation of *dog*. Because hetero-associative synapses are prominent in mediating the activation of word assemblies, we looked for the serial order mechanism in this set of connections. We notice that the phonemes-to-word connections show variability in their synaptic strength; the variability is visible in the rows of the lower-right block of Fig.5*A*. Following the theoretical results by Knoblauch & Pulvermüller (2005), we hypothesized that the different strengths encode the order of each pre-lexical unit and support sequence recognition capacity in the model. Thus, we now investigate the architecture of the synaptic weights in the phonemes-to-word connections and whether it also contributes to the word recognition capacities of the dendritic networks.

To test the hypothesis that average synaptic weights encode the phoneme serial position, we assigned an index to each phoneme, matching its serial position in the word, and determined the average synaptic connection between the phoneme and the word assemblies. For this analysis we considered the five lexica requiring memory. Because the weights of the phoneme-to-word connections vary across words and even more across the lexica, we normalized the synaptic weights by the strongest phoneme-to-word connection in each word. Similarly, because words have different lengths, we centered all the samples to their uniqueness points. The schematic on top of Fig.6*A* illustrates the serial position of the phonemes relative to the UP of the word *golden* from the *Overlap* lexicon. Here, the phoneme *E* corresponds to the uniqueness point, and the other phonemes are ordered according to it. The lower panel shows the average synaptic weight of each phoneme-to-word connection (y-axis) corresponding to the phoneme position in the word (x-axis). The minimum on the y-axis corresponds to the initial synaptic strength of the connections (*W*_0_).

The plot in Fig.6*A* shows significant differences among the three phonemes before the UP. The strength of the projections from phoneme to word increases with the serial position of the phoneme in the word: assemblies associated with early phonemes have weaker connections than those closer to the uniqueness point. In the present work, we have not investigated the dynamics leading to the formation of the weight architecture in Fig.6*A*. However, in the following, we show that this structure in the synaptic weights is necessary for recognizing words with phonological overlap, and it is not a spurious property of the network.

To test whether the increase of synaptic strength with serial order contributes to word recognition, we manipulated the network weights. We flattened the phoneme-to-words connections such that the curve in Fig.6*A* would appear flat (Methods). The panels in Fig.6*B* show a schematic of the transformation (top) and the resulting effective connectivity matrix (bottom) for the asymmetric Tripod network tested on the *Overlap* lexicon. For each post-synaptic word (y-axis), the strength of all incoming connections from the phoneme assemblies (x-axis) was identical. We then tested recognition accuracy as before and compared the two conditions (Fig.6*C*). We found that the structure of phoneme-to-word connections matters for the lexica that require memory but not for the other two (*Identity* and *No overlap*. Crucially, the *Digits* and *TIMIT* lexicon require memory, but they are less affected by the flattening of connections. The explanation is that if the words in the lexicon are distinguishable by the specific combination of phonemes, they do not rely on the serial order. The weight structure does matter when the phonological overlap is large, and the serial order is necessary to distinguish among words. Recognition accuracy drops for all dendritic models, losing 40 % to 60 % for the three lexical items with the highest degree of phonological overlap (i.e., *Overlap, Cohort, TISK*).

Further insights into the role of weight differences are obtained by comparing the average dendritic potential of words’ assemblies in the original and flattened conditions. The four panels in Fig.6*D* portray four words of the *Overlap* lexicon (*doll, poll, god,* and *dog*). The black line refers to the original model, and the orange line refers to the one with flattened phonemes to word connectivity; the values indicate the recognition score of the specific word. A zoom-in in the assembly dynamics is offered for the word *doll*. Fig.6*E* shows the membrane potential (top) and somatic activity (bottom) for all the neurons in the assembly in both the original network and flattened conditions; the traces are obtained averaging 10 presentations of the word. In the upper panels, the color scale is chosen such that only the NMDA spikes are visible (black reveals dendritic potential larger than 20 mV). Taken together, Fig.6*D* and *E* indicate that tampering with the effective connectivity matrix affects the average membrane dynamics of the word’s assembly. In the flattened condition, the membrane potential increases upon the presentation of early phonemes: between 0 ms to 100 ms, the assembly activity resembles the one in the original network. Then, at the presentation of the third phoneme (100 ms to 150 ms), the assembly fails to ignite the NMDA spikes that trigger the full reactivation. The result is that the membrane potential decays faster (orange traces), and the word population has sparser firing activity. Crucially, this does not entail that the word cannot be recognized at all, nor that all the words are equally affected, as evidenced by the word scores in Fig.6*H*. Rather, the weight structure offers an additional network mechanism that supports the distinction of words with full phonological overlap.

The present results illuminate pivotal role of dendrites in forming a structured and functional connectivity matrix. Models with dendrites support the formation of hetero-associative connections (phonemes to words), which are necessary for word recognition. The differences in the connection weights, which encode both the identity and order of the phonemes, are read by the activity of the phonemes assemblies and maintained in the dendritic memory of the word’s population. We now shift the focus from network structure to the short-term memory mechanism and show that it results from the interplay of non-linear dendritic integration and dendritic inhibition.

### Dendritic non-linearity governs temporal integration

The results have shown that networks of dendritic neurons are better at achieving word recognition than networks of point neurons. The sequence detection capacity follows from delayed temporal integration and feedforward connectivity between phonemes and word assemblies. We evidenced how dendritic memory in segregated compartments is the mechanism supporting the computation in dendritic networks. However, to activate and maintain the dendritic memory, the cells must be at a specific operational point that allows the expression of NMDA spikes. In addition, to avoid encoding the wrong memory, the network has to control the dendritic non-linearity and suppress NMDA spikes upon spurious activity in the assemblies. The voltage dependency in the NMDARs and tight dendritic inhibition permit such a computational state. We now show that the interactions of these mechanisms are necessary for the network model to achieve word recognition.

In the Tripod network, the dendritic non-linearity is governed by the ratio of NMDA-to-AMPA peak-conductances of the glutamatergic receptors (NAR) and the decay timescale of the NMDAR receptors (*τ_d_*). Fig.7*A* illustrates the impact of variations in the NMDA receptor on the dendritic membrane potential of the post-synaptic cell. The panels show the excitatory post-synaptic potential (EPSP) of a 300 µm dendrite following an excitatory spike on a synapse with weight 100 nF. In the left panel, the NAR varies between 0 and 2.7; in the right panel, the decay timescale *τ_d_* ranges from 2 ms to 50 ms. In both cases, the non-varied parameter maintains the baseline value of the model, NAR = 1.8 and *τ_d_* = 34 ms. The dashed lines portray this couple of parameters. The curves in the left panel show a NAR threshold for the onset of the non-linear response, approximately at NAR = 1.2. Above this NAR, the dendritic membrane enters a long-lasting depolarized state called *plateau potential*. These plateau potentials can be viewed as a form of intra-cellular dendritic memory on short timescales, and the conditions that elicit such states in the Tripod neuron have been described in Quaresima *et al*. (2022). The right panel in Fig.7*A* shows that the duration of plateau potentials was proportional to the timescale of the NMDA decay, *τ_d_*.

Inhibitory control is determined by the potential *V*_0_ that sets the target value for the voltage-dependent iSTDP rule on the dendrites of I2neurons. To determine the role of inhibitory plasticity v-iSTDP on synaptic learning, we varied the dendritic target voltage *V*_0_ in the range of −90 mV to −40 mV. The histograms in Fig.7*B* show the distribution of the post-synaptic weights from I2 neurons onto the dendritic compartment after the association phase. For a target value of *V*_0_ = 70 mV or lower, the synaptic weights accumulated at the maximum value 243 pF that we allowed for inhibitory synapses. When the target *V*_0_ was set higher, inhibitory synapses were potentiated less during the association phase, and the weight distributions flattened towards smaller values.

Changes in the dendritic non-linearity and inhibitory control do not imply that the network cannot process the phonemic stimuli correctly. Indeed, because the connectivity matrix remains unchanged, it may still drive correct assembly reactivation against weaker non-linearity or higher dendritic noise. In the present study, the structure of the excitatory connections is frozen to the one obtained from the associative phase. Thus, it could be that the parameter changes do not affect the task performance but only change the network dynamics. We measured the word recognition performance on all the parameter ranges to test this hypothesis. The three panels in Fig.7*D* illustrate the performance loss corresponding to networks simulated with each parameter variation. The score obtained for baseline values is used as the comparison term, and the loss is the difference between the score in the test condition and the score achieved in the baseline (*κ*-score ≈ 0.85). From the comparison of the three parameters swaps, it results that the inhibitory control plays the most significative role, with performance dropping of 60 % when the *V*_0_ is increased to −55 mV. To match the same loss, the NAR has to diminish to 0 to 0.6 and the timescale to 2 ms to 10 ms, which corresponds to absent plateau potentials.

Variations in the NMDA non-linearity and the strength of inhibitory control can also be traced in network activity. To investigate changes in the dynamics, we compared the average dendritic membrane potential of a word population with the average in the rest of the network while presenting the associated phoneme sequence. The difference between the control and the dendritic potential of the target population (*poll*) is shown in Fig.7*C* for the variations in the NMDARs and dendritic inhibition parameters. The left and middle panels show similar trends in that the difference in membrane potential stays close to zero for weak and short dendritic non-linearity. The activity of the target word assembly peaks after the first phoneme (*P*) but then returns to that of the control condition. On the other hand, if the non-linearity is parameterized as in the baseline model (black line), the dendritic membrane potential increases throughout the presentation of the phoneme sequence until it reaches a peak around the word offset and. A different pattern can be observed when the target potential *V*_0_ of v-iSTDP is varied. The rightmost panel in Fig.7*C* shows that the ERP is wider when dendritic inhibition is strong, although weaker inhibition also generates a substantial ERP relative to control. When *V*_0_ is equal to −70 mV or lower, the difference in membrane potential is nearly identical because the I2 inhibitory synapses that have developed during the association phase are similar (see Fig.7*B*). The changes in the network activity expressed by the difference in membrane potential (Fig.7*D*) match the *κ*-score loss of the word recognition task match (Fig.7*C*). This indicates that the dynamics portrayed in Fig.7*D* are causal for the task.

Crucially, in the latter case, the difference in membrane potential has a direction that seems counter-intuitive regarding the inhibitory function; more potent dendritic inhibition should cause weaker dendritic depolarizations. Hence, we inspected the average membrane potential for the word assembly and control condition to shed light on the ongoing dynamics. The four panels in SI Fig.9 show the soma and dendritic average potentials and solve the paradox of the role of v-iSTDP control. When the inhibitory target potential is larger, e.g., *V*_0_ =−50 mV, the network dendrites are permanently depolarized, and the difference between the target population activity and the control condition are less significant. Thus, the dendritic inhibition acts as a signal-to-noise control mechanism that allows the assembly to reactivate only upon the correct stimulus, maintaining the cells hyperpolarized otherwise.

We reasoned that the loss in recognition could be caused by testing recall with parameters different from those used in the associative phase, hence the mismatch. Thus, to exclude this possibility, we modulated dendritic non-linearity and inhibitory control during the associative phase and again tested word recognition. The panels in Fig.8*A, B* show the *κ*-score for a range of NARs, *τ_s_*, and *V*_0_ and indicate that enabling STDP plasticity does not solve the issue. Recognition drops for NAR below 1.0, for timescales shorter than 10 ms, and inhibitory target potential (*V*_0_) larger than −60 mV. In addition, there is an interaction between the NMDAR timescale and synaptic efficacy (NAR); for weaker non-linearity, longer timescales are beneficial. Rather than stabilizing, changes in the dendritic non-linearity and inhibitory control that occur during the associative phase also affect the formation of recurrent and hetero-associative connections. The average strength of the recurrent connections diminishes for most of the parameter variations and so does for the phonemes-to-word, in both cases the trends are similar to the recognition score (Fig.8*C, D*). These latter results indicate that both dendritic non-linearity and inhibitory control are necessary for the dendritic network to form assemblies. In the case of the dendritic linearity, this can be explained by the high threshold for synaptic strengthening in the STDP. The threshold is at −20 mV and if the dendrites are not sufficiently depolarized they will not enter the potentiation range. For the inhibitory control, the mechanism is more subtle. The strong external stimuli will override any network activity. Consistently with this observation, recurrent associations form despite weaker inhibitory control (Fig.8*D*). Conversely, the hetero-associative connections seem to require a high signal-to-noise ratio in the assemblies, and when inhibitory control weakens, this class of connections rapidly fades (Fig.8*F*).

**Figure 8:**
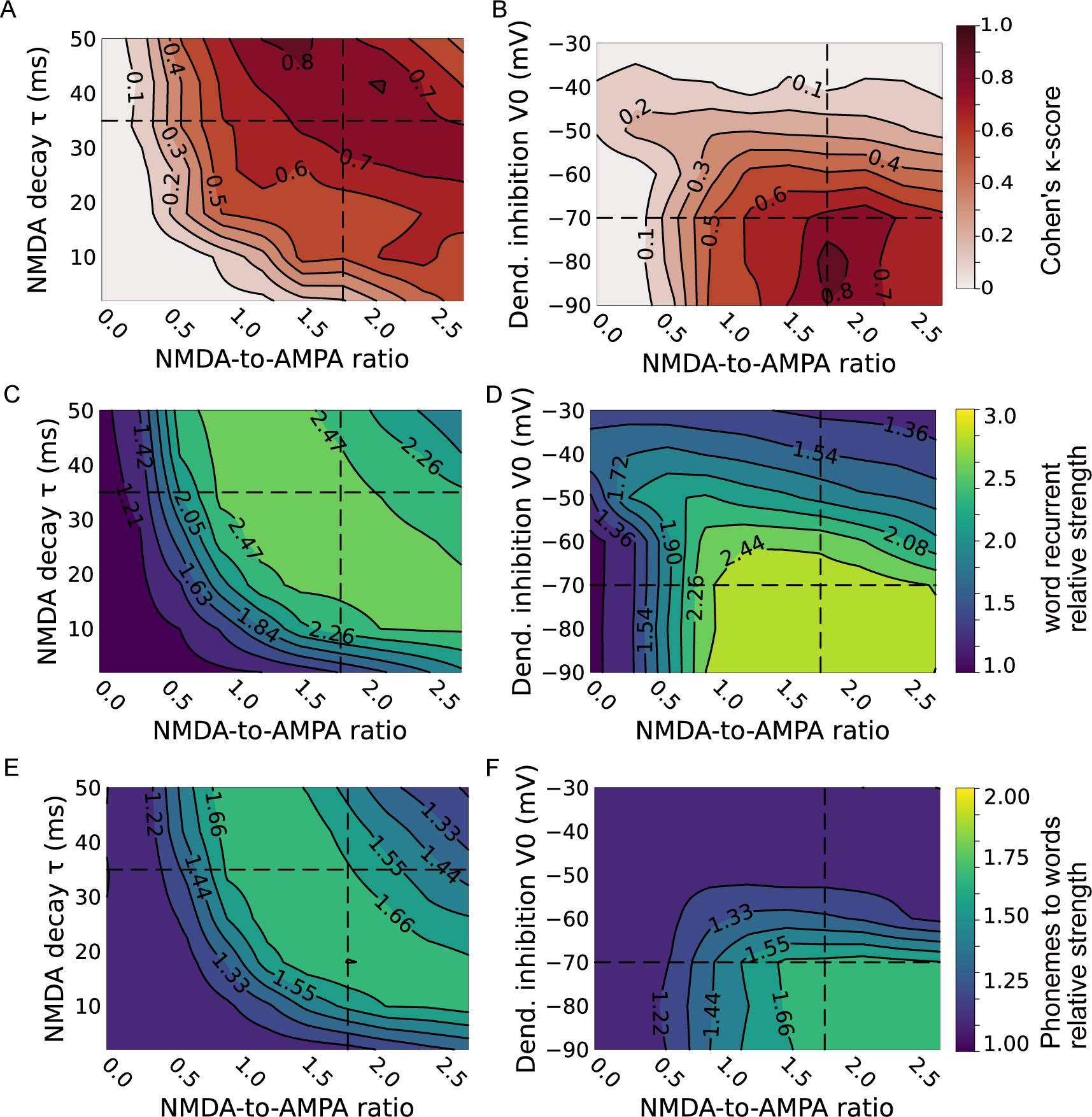
Dendritic non-linearity and inhibitory control enable the formation of phonemes to word connections. (**A**, **B**) Cohen’s *κ*-score of the Tripod asymmetric model for pairs of NMDA-to-AMPA ratio (NAR) and *τ_d_* and for pairs of NARs and *V*_0_. The model’s parameters are varied in both the associative and recall phases. The dashed lines indicate the baseline model. The left panel indicates that the *κ*-score is stable if changes in the NMDAR’s timescale are compensated with the NAR. Conversely, the interaction between the NAR and *V*_0_ variables is weaker (right panel); the performances decay linearly on both axes. (**C**, **D**) Strength of the recurrent word connections for the same parameter swap. Connections’ strength is divided by the average synaptic weight (10 pF) to evidence the formation of engrams. The contour plot of the words-to-words connection mirrors the *κ*-score panels, indicating that the changes in the dendritic non-linearity and inhibitory control also limit the formation of hetero-associative connections. (**E**, **F**) As in previous panels, for the phonemes to word connections. The hetero-associative connections respond similarly to the auto-associative for variations in the dendritic non-linearity (panel *E*). Conversely, weaker dendritic inhibition is more detrimental for the phonemes-to-word connections than for the recurrent ones.

To conclude, we have shown that dendritic non-linear excitability is necessary for the network’s word recognition capacity. The contribution of dendritic inhibition is also crucial because it governs dendritic depolarization and thus reduces the possibility that NMDA spikes originate from the intrinsic fluctuations in the network activity. The analysis suggests that NMDA spikes and tight inhibitory control are necessary during both the recall and associative phases. In the former, they contribute to forming the phonemes-to-word connections; in the latter, they allow for the temporal integration that is necessary for word recognition.

## Discussion

The present work investigated the formation and reactivation of Hebbian cell assemblies by stimuli with temporal structure. To this scope, we propose a novel, biologically constrained, spiking neural network model with dendrites and plastic excitatory connections. We evaluated the model in a phonemes-word association task and compared it with control networks with and without dendrites. The networks received sparse and overlapping excitatory projections, representing words and phonemes. The task was to activate the correct word assembly following the phonemic sequence. We demonstrated that the introduction of segregated dendritic compartments endowed the network with the capability to recognize sequences of inputs with overlapping features. In dendritic models, paired stimulation and STDP organized the networks in auto-associative and hetero-associative engrams. Word memories were then correctly recollected in vocabularies with partial or complete phonological overlap (as in the words *dog* and *god*). In contrast, the point-neuron models tested failed in the word recognition task when the vocabulary contained words with shared phonemes and did not form hetero-associative connections. In the following, we first discuss the significance of the present result for theories of biological memory, then we outline the contribution of dendritic non-linearity and inhibition in memory access.

The perception and recognition of phonological sequences is a fundamental human cognitive capacity (Povel & Essens, 1985; Dehaene *et al*., 2015). The recollection of sequence memories entails that the neural processes transform temporal patterns of activity into spatially-coded and time-compressed representations of the stimulus (Bagur *et al*., 2022; Fox *et al*., 2020; Chan *et al*., 2014; Vaz *et al*., 2020). However, how such a computation is carried out in biological networks remains poorly understood. First, it is not clear how word memories are stored in the neural substrate through Hebbian plasticity (Poeppel & Idsardi, 2022). Second, it is unknown how the networks integrate the short-term memory of the acoustic stimuli with the long-term memory of the lexical item to be accessed (Norris, 2017).

Concerning the storage of long-term memories, the dominant hypothesis is that consolidated memories are maintained in the strong recurrent connections of cell assemblies (Pulvermuller, 1999; Amit, 1995; Poo *et al*., 2016; Fuster, 1997). Cell assemblies and synaptic engrams successfully account for associative memories and their principles have been implemented in several computational models. Some of these studies have shown that associative plasticity, such as spike-time dependent plasticity (STDP), supports the acquisition of long-term synaptic memories and their recollection (Litwin-Kumar & Doiron, 2014; Zenke *et al*., 2015; Tomasello *et al*., 2018; Garagnani *et al*., 2009). However, these forms of associative memories seems to fall short in encoding relationships (Gallistel, 2021), such as the sequential order of phonemes in word memories. Our Tripod model shows that introducing dendritic compartments resolves this conundrum and supports the formation of order-sensitive memories.

The Tripod network model acquired word memories via STDP and maintained the memories in the hetero-associative connections between phonemes and word assemblies. The network memories were sensitive to both the identity and the order of the phonemes presented in the input. Information about the order was stored in the synaptic connections, and the strength depended on the serial position of the phoneme in the word. Weights were larger for phonemes closer to the uniqueness point. A similar mechanism for sequence detection in cell assemblies was already proposed by Knoblauch & Pulvermüller (2005). In contrast, the two point-neuron models investigated did not form hetero-associative connections for vocabularies with phonological overlap, let alone the sequence detection synaptic architecture. Remarkably, the DM model from Duarte & Morrison (2019) did better than the LKD model (Litwin-Kumar & Doiron, 2014). The main difference between the two models was related to the presence of longer timescales in the somatic NMDA receptors in the DM model. Similar to the Tripod network, the presence of slow-decaying depolarizing currents contributes significantly to the formation of the phonemes-to-words connections. The fact that these connections form in the case of memory-less vocabularies rules out that the models fail because of a shortage of synaptic budget in the face of the homeostatic mechanism.

The presence of hetero-associative connections is, however, not sufficient for recollecting the word memories. The model requires that the dendritic compartments express NMDA spikes. The nonlinearity in the segregated dendrites of the Tripod neuron mediates the interaction of short (dendritic) and long (synaptic) memories. Upon the presentation of external phonemic stimuli, the synaptic variables are read and encoded into the dendritic plateau potential of single cells. The integration of successive pieces of information is expressed in the slow build-up of the assembly dendritic membrane potential. The dendritic memory allowed for the integration of these sources of information over the word’s timescale. Crucially, the synaptic and dendritic memories have different computational trade-offs in terms of stability, robustness to noise, and duration of their transients (Chaudhuri & Fiete, 2016). The dendritic memory is encoded within a few tens of milliseconds and erased within hundreds while the engrams form over tens of seconds and remain stable over time (Appendix B). While stable memories have been shown to form in point-neuron models Litwin-Kumar & Doiron (2014), the novelty of our results is that also hetero-associative connections are stable and can re-activate overlapping memories.

To understand the relevance of the present contribution we must clarify how the word recognition task presented distinguishes from other instances of sequence learning. Sequence memories can also be instantiated as a chain of neural assemblies that activate in fixed order (Almeida-Filho *et al*., 2014, Hebb’s phase sequences). This neural phenomenon is observed during hippocampal replay and in bird song (Buzsáki, 2010) and can be reproduced in classical spiking network models (Fiete *et al*., 2010; Clopath *et al*., 2010; Gjorgjieva *et al*., 2011; Maes *et al*., 2020; Rajan *et al*., 2016; Reifenstein *et al*., 2021; Riquelme *et al*., 2023; Haga & Fukai, 2018; Gillett *et al*., 2020). In most cases, these models leverage hetero-associative connections between the chained assemblies. Nonetheless, the sequences investigated commonly include items that do not repeat. When there is overlap among the items, the model expresses additional features that allow distinguishing the identity of the sequence presented. For example, Cone & Shouval (2021) implement an external reservoir that depends on the identity of the sequence, while Maes *et al*. (2020) leverage an external neural clock.

Two elements set our model apart from these previous studies. First, the word memory is accessed at the word’s uniqueness point rather than at the end of the sequence. Early word memory access, at the uniqueness point, indicates that the memories are retrieved when sufficient information is in the input. Second, the sequence overlaps for long intervals, which impose temporal integration on the scale of a hundred milliseconds. Few studies have directly investigated whether STDP is sufficient for learning and retrieving word-like sequences. Among these, Duarte & Morrison (2014) investigated a point-neuron model with biological constraints and reported no net contribution of glutamatergic plasticity in sequence detection. Similarly, our attempts to implement sequence detection in the two control point neuron networks were not successful.

A few more recent studies have included dendritic computations in network models with weak physiological constraints (Hawkins & Ahmad, 2016; Leugering *et al*., 2023; Bouhadjar *et al*., 2022). Similarly to our model, these networks leverage plateau potentials to implement sequence detection. However, they present substantial abstractions in the dynamics of dendritic integration and synaptic plasticity. It is now well-known that dendritic processes determine the transfer function of single cells (Koch, 1998; Poirazi *et al*., 2003; London & Häusser, 2005; Ujfalussy *et al*., 2018; Payeur *et al*., 2019; Poirazi & Papoutsi, 2020; Jones & Kording, 2021). Their contribution is remarkable in synaptic clustering, processing memory, expanded memory storage, signal filtering and processing (Spruston, 2008; Quaresima *et al*., 2022; London & Häusser, 2005; Cazé & Stimberg, 2021; Jones & Kording, 2020; Mel, 1992; Poirazi *et al*., 2003, Baronig & Legenstein 2023 (in press)). However, it is not yet well understood how these computational primitives interact with the network dynamics and whether explicit modeling of dendritic processes can enrich the computational capabilities of neural networks (Larkum, 2022). Thus, the present work contributes to clarifying the computational implications of considering dendrites in realistic biological networks.

Our study implements a reduced three-compartment dendritic model (the Tripod neuron, Quaresima *et al*., 2022), and studies the effect of the dendritic computations in networks. By comparing dendritic models with one single compartment and with symmetric synaptic connections, we deduced that dendritic memory is the most important ingredient of the Tripod neuron for the task at hand. Dendritic memory is retained in the long-lasting depolarization (plateau potential) following the onset of NMDA spikes. In our network model, dendritic memory is strictly necessary for word recognition in the recall phase and contributes to the formation of phonemes-to-word connections in the associative phase. We have not tested if the same computations could be achieved leveraging other types of short-term memories, such as synaptic short-term adaptation (Mongillo *et al*., 2008; Cone & Shouval, 2021). Conversely, we verified that the same task cannot be achieved via neuronal memory (Fitz *et al*., 2020). The point neuron models were endowed with such memories in the form of membrane and spike-threshold adaptation.

Along with short-term memory, the two dendritic compartments of the Tripod neuron offer segregated pathways for signal integration. Each dendritic compartment supports a synaptic cluster; thus, it is balanced by dendritic inhibition and can autonomously elicit an NMDA spike (Quaresima *et al*., 2022). This property reflects the spatial distribution of synaptic clusters along the dendrites (Larkum, 2022; Kastellakis & Poirazi, 2019; Poirazi *et al*., 2003). Synaptic clusters allow the neuron to take part in multiple engrams because memories are maintained and reactivated independently (Legenstein & Maass, 2011; Kastellakis *et al*., 2016). We tested the implications of this capacity by comparing three dendritic models, of which only one model had two segregated dendritic pathways. Although their sequence-recognition performances were not significantly different, the asymmetric model required half of the specialized connections than the other dendritic models.

A third contribution of the dendritic compartments concerns the stability of the voltage-dependent plasticity rule. The segregated dendritic compartment spans approximately 60 mV between the hyperpolarized and depolarized state, and the potentiation region is accessible only to NMDA spikes. Thus, the dendritic non-linearity ensures that only salient stimuli are encoded in the long-term memory. Our results are in agreement with previous work on STDP in dendrites. These studies show that voltage-dependent STDP in the dendrites maintains stable memories across time (Kastellakis *et al*., 2016; Bono *et al*., 2017; Bono & Clopath, 2017) and that decoupling dendritic potential from somatic activity mitigates the stability-plasticity dilemma (Wilmes & Clopath, 2023).

The strengthening of excitatory connections makes the network prone to runaway excitation, a type of winner-takes-all dynamics (Ermentrout, 1992), and it destabilizes the network dynamics, causing continuous burst activity. In our model, the problem is solved by plasticity on the synapses of fast-spiking inhibitory neurons, which target the soma of tripod neurons and act fast compensatory mechanisms (Litwin-Kumar & Doiron, 2014; Zenke & Gerstner, 2017). However, somatic inhibition does not prevent dendrites from becoming strongly depolarized. This interferes with the formation of synaptic engrams. Following the experimental evidence on the role of inhibitory control on dendrites, (Herstel & Wierenga, 2021; Zucca *et al*., 2017; Chiu *et al*., 2018), we introduced dendritic inhibition as an additional compensatory mechanism, which can selectively silence dendritic branches, prevent the overriding of synaptic memories, and overall change the neuronal transfer function. The computational advantages of tight inhibitory balance on dendritic compartments were already reported by Yang *et al*. (2016) and Mikulasch *et al*. (2020).

The present work introduces voltage-dependent iSTDP in gabaergic synapses targeting dendritic compartments. The rule steers the dendritic compartment toward a hyperpolarized state. When the dendritic compartment is depolarized, the inhibitory synapses are strengthened. Our results indicate that this form of inhibitory control is necessary for sequence detection and for the formation of hetero-associative connections. However, our implementation of the voltage-based dendritic iSTDP presents some limitations. Our model requires strong dendritic inhibition, resulting in the inhibitory plasticity driving all the weights to the maximum synaptic efficacy, so there is no neuronal specificity in the inhibitory connections. We observed that lowering dendritic inhibitory control results in heterogeneous inhibitory synaptic weights but also reduces the recognition score. Future work should explore the inhibitory plasticity mechanism and individuate the correct parameters to develop neuron or assembly-specific inhibitory connections. This could be done by changing the plasticity rule with a non-spike-time-dependent one (Pedrosa & Clopath, 2020; Miehl & Gjorgjieva, 2022) or changing the connection probability to have more gabaergic synaptic contacts per dendrite.

Moreover, the lack of neuron-specificity in the inhibitory weights indicates that dendritic inhibition does not act as a competition mechanism. This is in contrast with other models of sequence processing (Maes *et al*., 2020; Cone & Shouval, 2021, Vlachos et al., in preparation) or word recognition (McClelland & Elman, 1986; Hannagan *et al*., 2013), in which inhibition mediates competition among the excitatory assemblies. Somatic inhibition could still play this role because its synaptic strength depends on the neuron’s firing rate (Appendix A). However, the lack of plasticity in the glutamatergic connections of the inhibitory cells (*E*→*I1*) makes it such that inhibition acts as a uniform blanket rather than a competition mechanism. Thus, our network solves the word recognition task without leveraging lateral inhibition among lexical competitors.

The homeostatic mechanism in the vSTDP, which maintains the incoming synaptic strength fixed for each neuron, is the only mechanism that can mediate competition, and its effect is expressed in the weak reciprocal connections among words. However, homeostasis seems ineffective. The connectivity matrices in the results and the appendices indicate that the assemblies that share some phonemes have, on average, stronger excitatory interactions than those that do not. We deem that implementing an explicit competition mechanism would increase the capacity of the network to distinguish among words with shared phonemes.

## Conclusions and future work

The present study proposed a highly biologically constrained model for spoken word recognition. We show that dendritic processes endow the network with the necessary computational primitives for storing and recalling word memories. Beyond the limitations hitherto discussed, we think three main improvements can be made to the model. in order to account for human word recognition. First, word populations should more actively compete among themselves. Blanket feedback inhibition works, but words with overlapping features tend to associate despite their mutual contribution, which is harmful to recognition. Anti-Hebbian plasticity implemented between inhibitory and excitatory neurons could introduce this active competition (Luz & Shamir, 2012) and carve part of the lexical memory in the inhibitory connections. Second, word recognition fails more often for long words. This is because the dendritic memory is relatively short-range, and the synaptic connections can effectively distinguish solely the order of three phonemes. Thus we think that introducing intermediate representations, such as syllables, may help sharpen the recollection of word memories. Third, to introduce feedback in the network, such that lexical decisions are made based on joined phonetic and semantic information. Future work should investigate the pathways through which contextual information can be delivered to the network (feedback pathways on the apical dendrites, disinhibitory circuits, etc. (Naumann *et al*., 2022)) and how this will affect the network dynamics.

## Methods

In this study, we examine the activity of spiking network models when exposed to external stimuli representing phonemes and words. In the following sections, we first describe the equations characterizing the neuron models and their synaptic dynamics. Then we describe the network architecture, the recurrent connectivity, and the plasticity rules for each network class. Finally, we describe the experimental protocol used to model word recognition.

### Neurons and synapses

The network models investigated are composed of two types of neurons, excitatory and inhibitory. The excitatory neurons have been modeled as three-compartment neurons in the Tripod networks and single-compartment units in the point-neuron networks. The inhibitory neurons have two subtypes, both modeled as single-compartment neurons. The models and parameters used vary for each network, and the variations are constrained to ensure stability during the simulation. For the two previously published networks, namely the L2/3 cortical circuit by Duarte & Morrison (2019) and the balanced network by Litwin-Kumar & Doiron (2014), we used models and parameters identical with the original publications; except for the plasticity rules introduced in the Duarte & Morrison (2019). In the following particular attention is reserved to the dendritic network, as it is the most complex model and constitutes the novel contribution of the present work.

### Excitatory neurons

#### Tripod neurons

The Tripod neurons comprise three electronically segregated compartments: an axosomatic compartment and two membrane patches. A detailed description of the model is presented in Quaresima *et al*. (2022), which also constitutes the second chapter of the present manuscript. The somatic compartment is modeled with the adaptive exponential integrate-and-fire model (Brette & Gerstner, 2005), it is two-dimensional and comprises the dynamics of membrane potential and adaptive hyper-polarizing currents (Eqs.1,2).

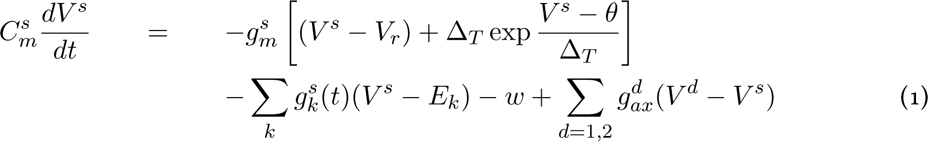

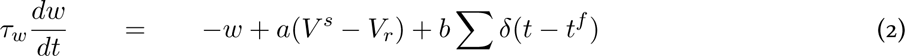

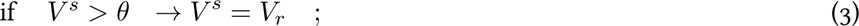

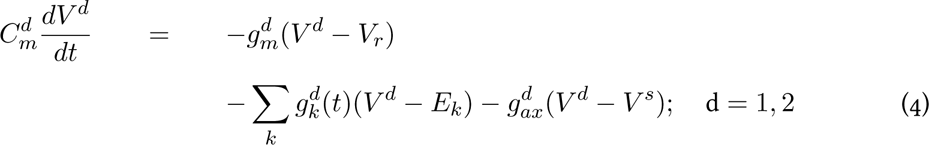

The leak conductance 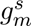 defines the electrical permeability of the somatic membrane, 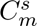 its capacitance, and 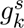 the set of variable synaptic conductances. The membrane potential *V ^s^* is reset to *V_r_* after a spike, and the adaptation current *w* is increased by a constant value *b* (Eq.2). Spikes occur at times *t^f^* when the potential *V ^s^* exceeds a threshold *θ* (Eq.3). Parameters for the somatic compartment were fixed and set to the values used in Brette & Gerstner (2005) and are listed in Table 1.

**Table 1:**
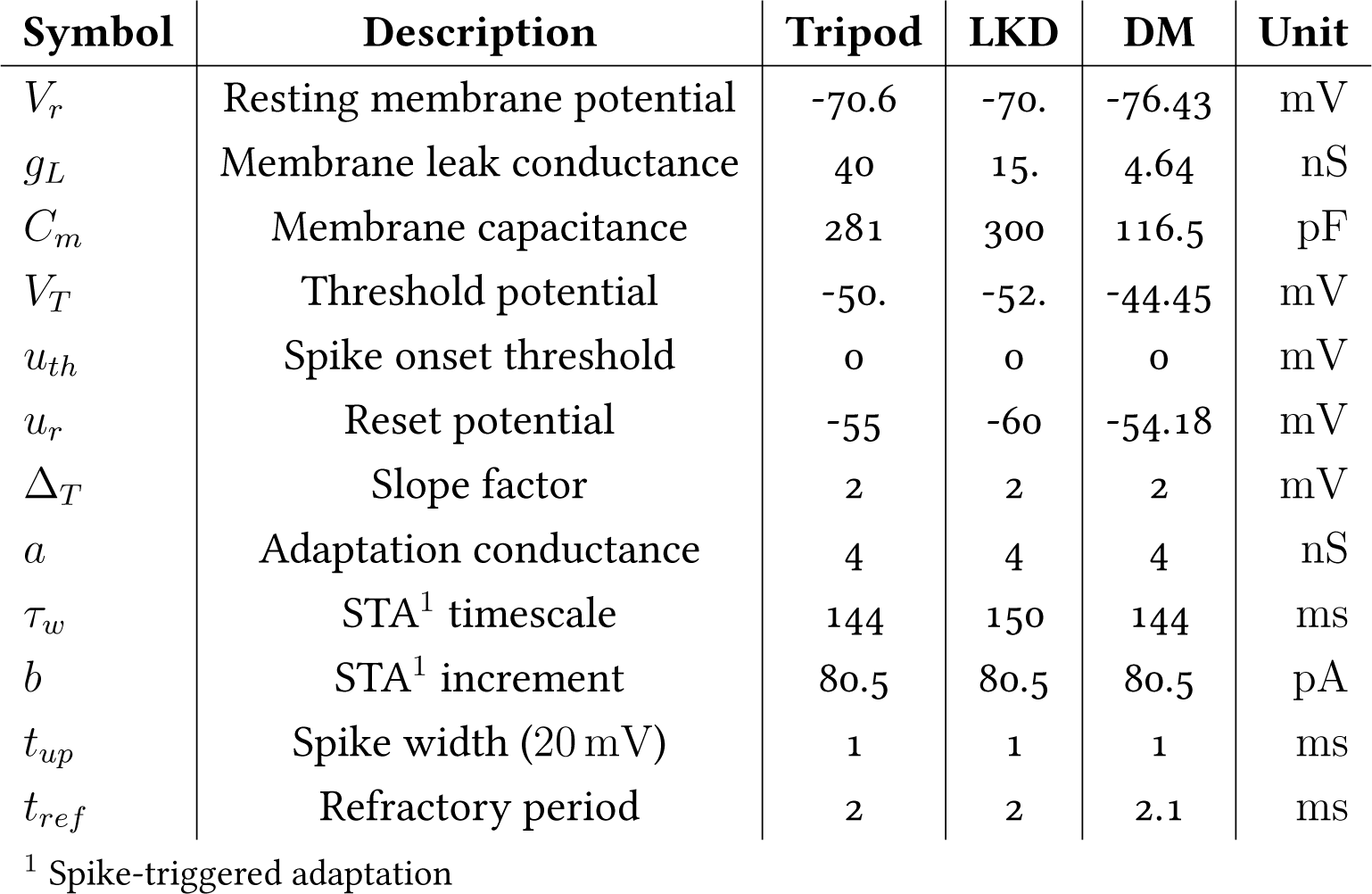
Parameters for the axosomatic compartment (Quaresima *et al*., 2022; Litwin-Kumar & Doiron, 2014; Duarte & Morrison, 2019).

The somatic compartment is coupled to the dendrites by the term 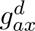 (Eq.1), it accounts for the axial conductance of the soma and the dendrite. The electrical properties of the dendritic compartment are governed by the passive membrane-patch equation Eq.4 (Koch, 1998). The equation’s parameters depend on the dendritic geometry combined with physiological parameters. The physiology concerns the membrane permeability, the membrane capacitance, and the axial impedance to the soma 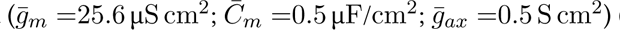 (Koch, 1998; Eyal *et al*., 2016). In the present study, we use the parameters obtained from human pyramidal cells Eyal *et al*. (2016) because they show enhanced dendritic memory (Quaresima *et al*., 2022). The dendritic diameter is fixed at 4 mm, and dendritic electrical properties are determined only by the dendritic length *L_d_*, in agreement with the equations:

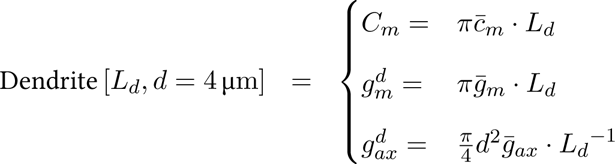

In all the dendritic models tested in the present study, the dendritic compartments have heterogeneous lengths in the range 150 µm to 400 µm.

The current flow between soma and dendrites depends on *g_ax_* (*V_d_ > V_s_*). It usually is positive (dromic); conversely, during somatic firing, the soma compartment’s membrane potential is clamped to 20 mV for the duration of the spike (spike width, 1 ms). During the spike interval, the current flow is antidromic (*V_d_ < V_s_*), and the current flowing to the dendrites acts as a backpropagating axon potential (the dendritic and soma potentials are illustrated in Fig.1*A*, the inset shows the potentials’ dynamics during somatic firing).

The Tripod neurons express AMPA, NMDA, GABA_A_, and GABA_B_ receptors on the dendrites and only AMPA and GABA_A_ on the soma. The synaptic transmission is modeled with a double-exponential equation (Roth & van Rossum, 2009). The equation describes the rise and decay of the receptors’ conductance *g_k_*. The synaptic conductance *g_k_* is given by:

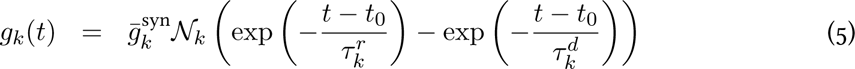

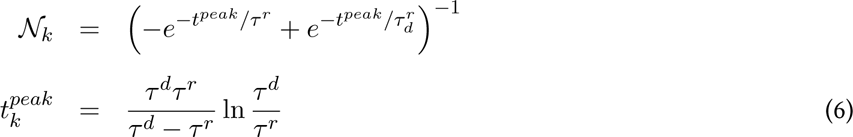

with *k* ∈ {AMPA, NMDA, GABA_A_, GABA_B_} indicating that each receptor has specific parameters. The timescales of rise and decay are given by *τ_r_* and *τ_d_* while the amplitude of the curve is defined by the maximal conductance parameter *g_syn_*. To ensure that the amplitude equals 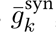, the conductance was scaled by the fixed normalization factor N*_k_* Eq.6. The ratio between the maximal conductance of NMDA to AMPA receptors (NAR) governs the non-linear response to glutamatergic stimuli, in the present work is set as in human-like cells in Eyal *et al*. (2018); Quaresima *et al*. (2022). The synaptic parameters of the Tripod neuron are listed in Table 3.

**Table 2:**
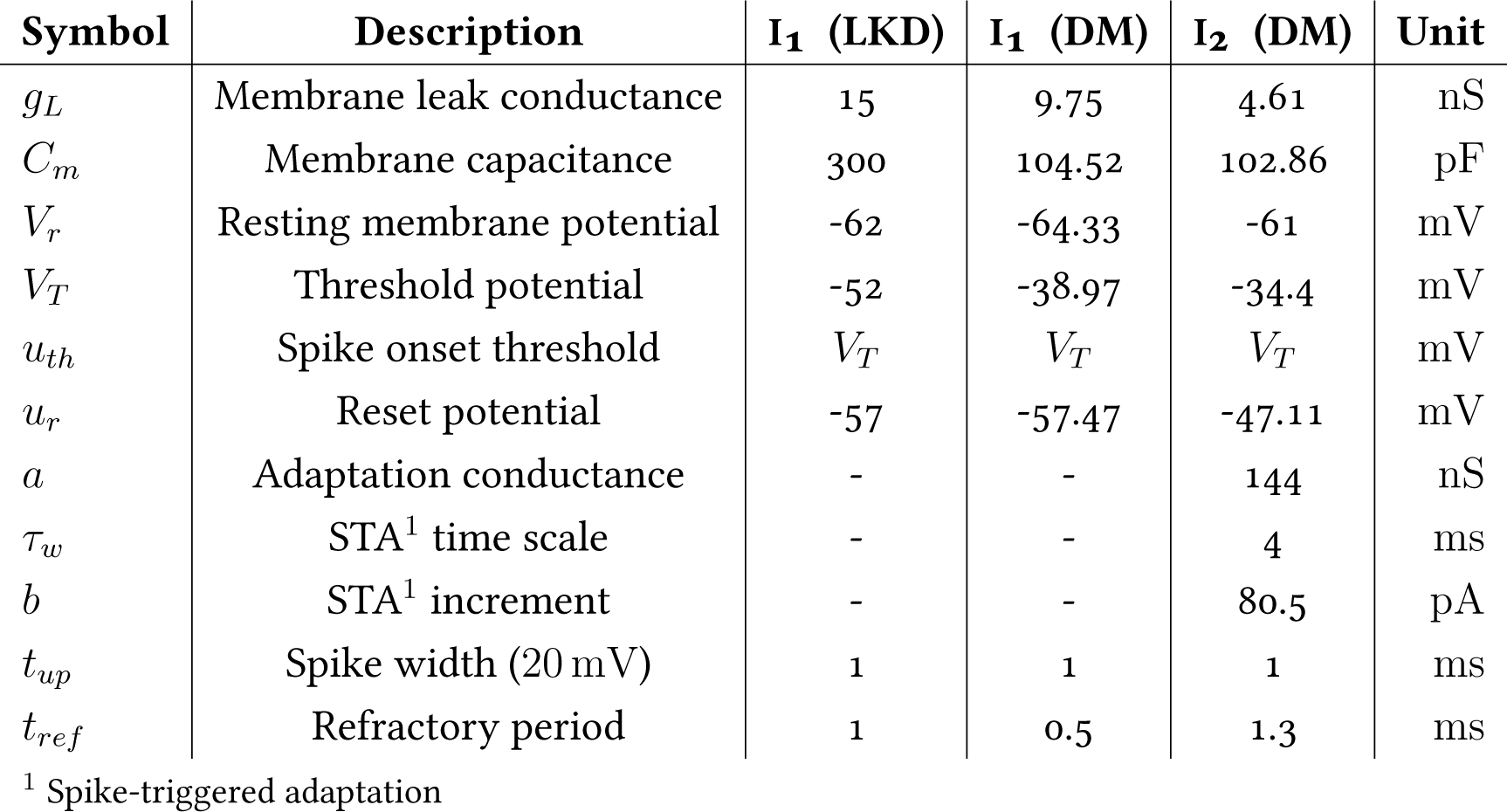
Parameters for the axosomatic compartment of the inhibitory neurons for the three network models (Litwin-Kumar & Doiron, 2014; Duarte & Morrison, 2019). The inhibitory neurons of the Tripod network have the same parameters as those in the Duarte and Morrison model.

| Symbol | Description | $I_1$ (LKD) | $I_1$ (DM) | $I_2$ (DM) | Unit |
| --- | --- | --- | --- | --- | --- |
| $g_L$ | Membrane leak conductance | 15 | 9.75 | 4.61 | nS |
| $C_m$ | Membrane capacitance | 300 | 104.52 | 102.86 | pF |
| $V_r$ | Resting membrane potential | -62 | -64.33 | -61 | mV |
| $V_T$ | Threshold potential | -52 | -38.97 | -34.4 | mV |
| $u_{th}$ | Spike onset threshold | $V_T$ | $V_T$ | $V_T$ | mV |
| $u_r$ | Reset potential | -57 | -57.47 | -47.11 | mV |
| $a$ | Adaptation conductance | - | - | 144 | nS |
| $\tau_w$ | STA<sup>1</sup> time scale | - | - | 4 | ms |
| $b$ | STA<sup>1</sup> increment | - | - | 80.5 | pA |
| $t_{up}$ | Spike width (20 mV) | 1 | 1 | 1 | ms |
| $t_{ref}$ | Refractory period | 1 | 0.5 | 1.3 | ms |

<sup>1</sup> Spike-triggered adaptation

**Table 3:** Synaptic parameters for the excitatory and inhibitory neurons in the Tripod network.

| **Receptor** | $E_r$ (mV) | $\tau_{rise}$ (ms) | $\tau_{decay}$ (ms) | $g_{syn}$ (nS) | $\gamma \left( \frac{1}{mV} \right)$ |
| --- | --- | --- | --- | --- | --- |
| **Tripod soma** |  |  |  |  |  |
| AMPA | 0.0 | 0.26 | 2. | 0.73 | - |
| GABA<sub>A</sub> | -70 | 0.1 | 15 | 0.38 | - |
| **Tripod dendrites** |  |  |  |  |  |
| AMPA | 0.0 | 0.26 | 2. | 0.73 | - |
| NMDA | 0.0 | 8. | 35. | $0.73 \cdot \text{NAR}$ | 0.075 |
| GABA<sub>A</sub> | -70 | 4.8 | 29.0 | 0.27 | - |
| GABA<sub>B</sub> | - 90 | 30 | 400 | 0.006 | - |
| **I<sub>1</sub> soma** |  |  |  |  |  |
| AMPA | 0.0 | 0.18 | 0.7 | 1.04 | - |
| GABA<sub>A</sub> | -75.0 | 0.19 | 2.5 | 0.84 | - |
| **I<sub>2</sub> soma** |  |  |  |  |  |
| AMPA | 0.0 | 0.18 | 1.8 | 0.56 | - |
| GABA<sub>A</sub> | -75.0 | 0.19 | 5.0 | 0.59 | - |

#### Point neuron models

The excitatory cells of the point-neuron networks are also implemented with the AdEx model (Eq.1). For the point-neuron models, the membrane potential threshold *θ* changes over time and increases after each spike. The dynamic threshold facilitates voltage-dependent synaptic plasticity during the bursting intervals, it is governed by the equation:

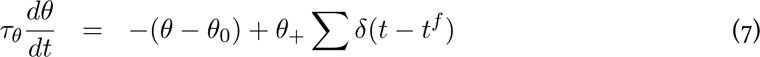

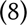

For the network borrowed from Duarte & Morrison (2019), the soma hosts four types of receptors, as in the Tripod. Conversely, the model from Litwin-Kumar & Doiron (2014) only has AMPA and GABA_A_ receptors. The parameters of the point-neuron AdEx models and synapses are listed in Table 1 and Table 3.

### Inhibitory interneurons

Inhibitory neurons are modeled as single, isopotential compartments with leaky integrate-and-fire dynamics. The membrane potential *V_i_* is governed by the following equations:

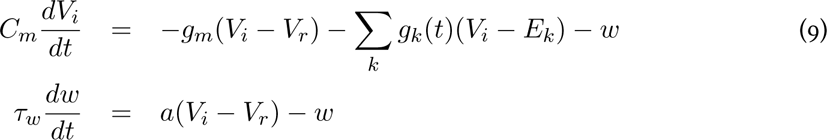

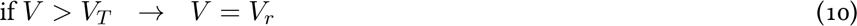

Following previous modeling efforts (e.g. Park & Geffen (2020); Yang *et al*. (2016); Duarte & Morrison (2019)) and consistently with physiological description (Tremblay *et al*., 2016), we implement two types of inhibitory interneurons: fast-spiking and non-fast-spiking. The first class corresponds to a set of well-defined parvalbumin-expressing (PV) GABAergic neurons, among which basket-cells; these cells primarily target the perisomatic regions and are indicated in this work as I1population. Conversely, the second group includes parvalbumin-negative GABAergic neurons expressing somatostatin. For notational simplicity, we refer to the second group as I2. Following Duarte & Morrison (2019), the fast-spiking neurons are modeled as not adaptive while the I2 neurons are. The two classes differ in LIF model parameters (Table 2), and synaptic properties (Table 3). In our model, they also differ for the compartment of the Tripod neuron they connect to. I1 neurons target the soma compartment, and their synapses express fast GABA_A_ receptors. I2 neurons, on the other hand, target the dendrites and, due to the additional slow GABA_B_ component of the dendritic synapses, have a longer timescale. The inhibitory neurons’ synapses express AMPA, GABA_A_ receptors, and the synaptic transmission is modeled with a double-exponential model. The parameters for the inhibitory neurons are borrowed from Duarte & Morrison (2019) and are reported in Table 1 and Table 4.

**Table 4:** Synaptic parameters for the excitatory and inhibitory neurons in the point neuron models (Duarte & Morrison, 2019; Litwin-Kumar & Doiron, 2014).

| Receptor | $E_r$(mV) | $\tau_{rise}$(ms) | $\tau_{decay}$(ms) | $g_{syn}$(nS) | $\gamma$ ($\frac{1}{mV}$) |
| --- | --- | --- | --- | --- | --- |
| **LKD soma** |  |  |  |  |  |
| AMPA | 0.0 | 1.0 | 6.0 | 1.0 | - |
| GABA<sub>A</sub> | -75.0 | 0.5 | 2.0 | 1.0 | - |
| **LKD I<sub>1</sub>** |  |  |  |  |  |
| AMPA | 0.0 | 1.0 | 6.0 | 1.0 | - |
| GABA<sub>A</sub> | -75.0 | 0.5 | 2.0 | 1.0 | - |
| **Duarte soma** |  |  |  |  |  |
| AMPA | 0.0 | 0.25 | 2.0 | 0.73 | - |
| NMDA | 0.0 | 0.99 | 100.0 | 0.16 | - |
| GABA<sub>A</sub> | -75.0 | 0.5 | 6.0 | 0.26 | - |
| GABA<sub>B</sub> | -90.0 | 30.0 | 100.0 | 0.01 | - |
| **Duarte I<sub>1</sub>** |  |  |  |  |  |
| AMPA | 0.0 | 0.1 | 0.7 | 1.6 | - |
| NMDA | 0.0 | 1.0 | 100.0 | 0.0 | - |
| GABA<sub>A</sub> | -75.0 | 0.1 | 2.5 | 1.0 | - |
| GABA<sub>B</sub> | -90.0 | 25.0 | 400.0 | 0.02 | - |
| **LKD I<sub>2</sub>** |  |  |  |  |  |
| AMPA | 0.0 | 0.2 | 1.8 | 0.8 | - |
| NMDA | 0.0 | 1.0 | 100.0 | 0.01 | - |
| GABA<sub>A</sub> | -75.0 | 0.2 | 5.0 | 0.7 | - |
| GABA<sub>B</sub> | -90.0 | 25.0 | 500.0 | 0.02 | - |

### Network architectures

We implement three main classes of networks that vary for the number and types of neurons considered and for the connectivity between them. The novel network model is composed of Tripod neurons as excitatory cells. The network counts 2000 Tripod neurons with dendritic lengths uniformly distributed in the range150 µm to 400 µm; along with 175 fast-spiking neurons, and 325 slow-spiking neurons. The proportion of inhibitory neurons follows Duarte & Morrison (2019). The network lacks geometrical structure and is conceived as a local cortical circuit. The Tripod neurons connect reciprocally through the dendrites and to the soma of inhibitory neurons. The fast-spiking neurons target the Tripod neurons’ soma, while the slow-spiking neurons target the dendrites. Inhibitory neurons also connect reciprocally. All connections have a probability of *p* =0.2. The connections targeting the two dendrites of a Tripod neuron are drawn independently from each other. Hence some neurons will connect through one dendrite only, and others will connect through both dendrites (pre-synaptic spikes arrive separately on both compartments). The connectivity pattern described is the default network configuration and is identified as the *asymmetrical* network. As a result, the Tripod network has 10 different connection types; the Tripod-Tripod and I2-Tripod are doubled because they account for the two dendrites.

The remaining two networks are composed of point-neuron models, illustrated in the right panel of Fig.1*B*. One of the models implements an exact copy of the network described in Litwin-Kumar & Doiron (2014) (LKD model). The network is composed of 4000 excitatory neurons and 1000 inhibitory neurons. Then neurons are sparsely connected with a probability, *p* =0.2. The third network is composed of 2000 excitatory neurons, 200 fast-spiking neurons and 200 slow-spiking neurons. The network is modeled after the non-heterogeneous model of cortical L2/3 (Duarte & Morrison, 2019). Neurons are connected with probability and connectivity strength that depends on the type of connection, for a total of nine types of connections. The network is dubbed the DM network.

The networks’ neuron types and the number of cells are listed in Table 6. In all the models, the synaptic weights change according to the connection types and are listed in Table 7 with the respective connection probability. An illustration of the Tripod and point neuron networks connectivity is offered in Fig.1*B*

### Synaptic learning rules

The synapses connecting excitatory neurons (EE) and from the inhibitory to the excitatory neurons (IE) are subject to spike-time-dependent plasticity (STDP). The two plasticity rules are illustrated in Fig.1*C*.

#### Excitatory STDP

The glutamatergic synapses undergo a voltage-dependent STDP rule (Clopath *et al*., 2010). The voltage-dependent STDP (vSTDP) strengthens the connection when, following a pre-synaptic spike, the post-synaptic neuron’s membrane potential is depolarized. The equations governing the vSTDP rule are:

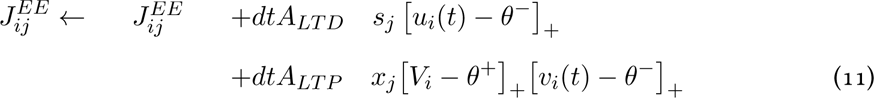

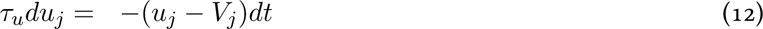

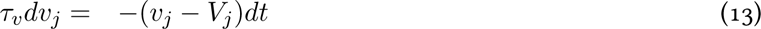

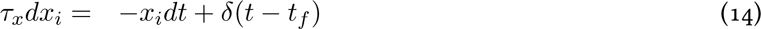

The 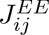 indicates the connection between the excitatory neurons, post-synaptic *i*, and pre-synaptic *j*. The variable 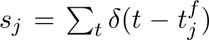 is the spike train of the presynaptic neuron, and *x_i_* is the filtered spike train of the post-synaptic cell. In addition, the variables *V_j_, u_j_, v_j_* are the membrane potential, the slow, and the fast-filtered membrane potential of the targeted compartment. The two thresholds (*θ*^+^*, θ^−^*) mark the region where long-term potentiation (LTP) and long-term depression (LTD) occur, coupled with the learning rates *A_LT P_* and *A_LT D_*. In the point-neuron models, the vSTDP is applied to the soma 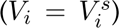, while in the Tripod neurons, the vSTDP is applied to the dendrites 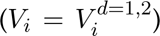. For each couple of connected Tripod neurons, there are two values 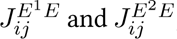 one for each dendrite, and they are updated independently. In addition, the vSTDP rule includes a homeostatic mechanism that maintains the sum of pre-synaptic weights fixed; the homeostatic rule follows the prescription of synaptic scaling (Turrigiano, 2011; Tetzlaff *et al*., 2011; Triesch *et al*., 2018) and is multiplicative. Conversely, the homeostatic rule in Litwin-Kumar & Doiron (2014) is additive. The multiplicative homeostatic rule is implemented as follows:

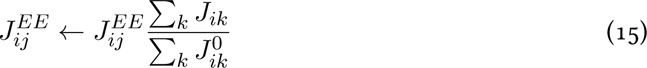

where 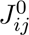 is the initial value of the synaptic weight. Homeostasis is applied every 20 ms.

#### Inhibitory STDP

For the connections from inhibitory to excitatory neurons, the networks implement the iSTDP rule from Vogels *et al*. (2011). The iSTDP governs the network stability by regulating the inhibitory synaptic strength. It is a necessary ingredient for the formation of assemblies in the original LKD network because it prevents winner-takes-all dynamics. We implemented the same mechanism for the I1or fast-spiking interneurons in the three networks, (*E^s^*I1); the iSTDP maintains the firing rate of the excitatory cells by modulating the synaptic strength of the inhibitory pathways. In addition, we modified the iSTDP rule for the slow interneuron type such that they can control the dendritic non-linearity of the Tripod’s dendrites. The I2iSTDP targets the membrane potential of the post-synaptic neuron, rather than its firing rate. We refer to it as voltage-dependent iSTDP or v-iSTDP. It applies to synapses on the soma (*E*[*s*]I2) in the DM and to synapses on the dendrites (*E*[*d*]I2) in the Tripod network. Because there are non I2neurons in the LKD network, this plasticity rule does not apply there. The two iSTDP rules follow the equations:

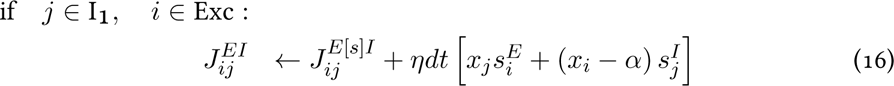

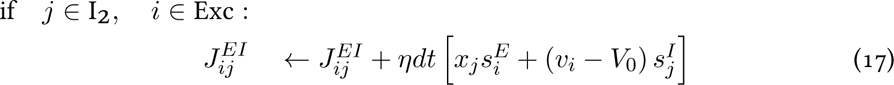

The 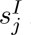 is the spike train of the inhibitory neuron *j*, *x_i_* is the filtered spike trains of the excitatory neuron *i* and *v_i_* is the dendritic or somatic potential (depending on the network to which the plasticity rule applies), respectively. The *α* is the target firing rate for the iSTDP of I1neurons, while the *V*_0_ is the target membrane potential for the iSTDP of I2neurons. *η* is the learning rate. The parameters of the iSTDP rules are listed in Table 5.

**Table 5:** Parameters for the v-STDP, iSTDP, and v-iSTDP plasticity rules. The v-STDP parameters for the point-neuron models (LKD and DM) are obtained from Clopath *et al*. (2010), and identical to the original study by Litwin-Kumar & Doiron (2014); for the dendritic model is obtained from Bono & Clopath (2017). The iSTDP and v-iSTDP rules follow the parameters by Vogels *et al*. (2011) with variations on the learning rate *η* and the target of the homeostasis for the two interneurons.

| Symbol | Description | LKD | DM | Tripod | Unit |
| --- | --- | --- | --- | --- | --- |
| **v-STDP** |  |  |  |  |  |
| $a^-$ | LTD learning rate | 8.0E-05 | 8.0E-05 | 4.0E-05 | Hz |
| $a^+$ | LTP learning rate | 1.4E-04 | 1.4E-04 | 1.4E-04 | Hz |
| $\theta^-$ | LTD threshold | -70 | -70 | -40 | mV |
| $\theta^+$ | LTP threshold | -49 | -49 | -20 | mV |
| $\tau_u$ | slow membrane filter | 10.0 | 10.0 | 15.0 | ms |
| $\tau_v$ | fast membrane filter | 7.0 | 7.0 | 45.0 | ms |
| $\tau_x$ | spike train filter | 15.0 | 15.0 | 20.0 | ms |
| $\tau_s$ | homeostasis timescale | 20.0 | 20.0 | 20.0 | ms |
| $j^-$ | minimum synaptic weight | 2.78 | 2.78 | 2.78 | pF |
| $j^+$ | maximum synaptic weight | 21.0 | 4.0 | 41.4 | pF |
| **iSTDP** |  |  |  |  |  |
| $\eta$ | learning rate | 1.0 | 0.2 | 0.2 | Hz |
| $j^+$ | maximum synaptic weight | 243.0 | 243.0 | 243.0 | pF |
| $j^-$ | minimum synaptic weight | 2.78 | 2.78 | 2.78 | pF |
| **I<sub>1</sub>neurons** |  |  |  |  |  |
| $r_0$ | target firing rate | 5 | 10 | 10 | Hz |
| $\tau_y$ | rate filter timescale | 20.0 | 20.0 | 20.0 | ms |
| **I<sub>2</sub>neurons** |  |  |  |  |  |
| $vd$ | target dend. potential | - | -70 | -70 | mV |
| $\tau_d$ | dend. potential filter timescale | - | 5.0 | 5.0 | ms |

### Stimuli

#### Phonemes and word projections

The networks were stimulated through external projections encoding phoneme and word inputs. Depending on the lexicon used, each network has 20 to 35 distinct projections, each representing a distinct phoneme or a word. Each projection targets *ρ* = 5 % of the network, creating an assembly of co-activated cells. The assemblies were approximately 100 neurons in the Tripod and DM network and 200 in the LKD network. The projections are distributed randomly to the network’s neurons; therefore, the assemblies lack any geometrical structure and each cell can be contained in more than one assembly. The phoneme and word assemblies were stimulated with spike trains. All pre-synaptic projections have fixed firing rate (8 kHz) and synaptic efficacy (2.78 pF for point-neuron models, 20 pF for the Tripod model). These numbers were optimized to have a strong neuronal response but not unstabilize the network. Stimuli spike trains are drawn from a Poisson distribution at every presentation, with the rate being the same for all projections; as a consequence, the stimuli cannot be interpreted as rate- or time-coded. The only information available to the networks to discriminate between the stimuli is the identity of the activated assemblies, which constitutes a spatial code. The phoneme stimuli were presented for 50 ms each, and the word stimuli were presented for the entire duration of the stimulus. Each word was followed by a silent interval without external stimuli, the pause lasts 50 ms.

#### Sequence association and recall

The stimulation protocol consists of two phases during which the phonemes to words associations are first established and then recall is tested, both phases last 5 min. During the associative phase, we present the network with external stimuli on both phonemes and word projections. The stimuli are presented in sequence; one word is drawn randomly, and the corresponding phoneme projections are activated. The phonemes populations are activated for 50 ms each, while the word ensembles are activated for the entire duration of the word (that is 50 ms times the number of phonemes contained in the word). The number of words presented in the two phases depends on the average length and the interval duration; for the experiments discussed here, each word is presented 50 to 100 times. All words are presented approximately the same number of times; therefore, there are no frequency effects. The phase duration has been chosen to be long enough for the synaptic weights to converge but short enough to allow for several simulations and maintain the study reproducible.

#### Lexica

The lexica were chosen from previous publications or assembled for this specific study. The seven lexica tested vary for the number of items, the average length of words, and the phonological overlap. Phonological overlap is computed as the ratio between the number of words in the lexicon and the number of phonemes: O = *N_w_/N_p_*. The lexica and their characteristics are listed in Table 8.

The TIMIT lexicon has been selected based on the work from Dong *et al*. (2018). The Cohort lexicon was obtained from Granger *et al*. (1994). The TISK lexicon was obtained from Hannagan *et al*. (2013). The remaining lexica are original to this work.

### Word recognition measure

#### Uniqueness and offset point

We individuate two salient time points for word recognition for every phoneme sequence presented during the recall phase. One is the offset of the word (offset point, OP), that is, the last phoneme of the sequence. The other is the word’s uniqueness point (UP). The UP corresponds to the phoneme that permits distinguishing a word from any other in the lexicon (Marslen-Wilson, 1973); when the word is contained in another word (such as *gold* and *golden*), the UP is the phoneme following the last shared phoneme, i.e., the silence after the end of the word in case of *gold*. As in the previous section, we estimate word recognition for every trial by measuring the firing rate in the adjacent interval. We associate the OP with an interval lasting the entire length of the phonemes’ sequence, from the word onset to the word offset (50 ms times the length of the word); this measure is the same one portrayed in Fig.2*D*. Conversely, the UP interval is shorter and corresponds to the duration of one single phoneme (50 ms); in this case, the interval starts and ends with the UP phoneme. An example of the uniqueness and offset points of the word *golden*, and their intervals, is illustrated in Fig.3*A*; the offset and UP time-points are marked on the phoneme sequence, respectively, by the blue and green arrow marks. Because words are longer than one phoneme, the interval associated with the offset is, on average, longer than the one associated with the UP.

#### Recognition *κ*-score

Because the reactivation of word assemblies is delayed with respect to the onset ofset of the phoneme sequence, we introduced a variable time shift in the measure of the firing rate to individuate the optimal interval of word recognition. For both the offset and UP measures, the measured intervals were probed with delays from −150 ms to 200 ms, sampling every 5 ms. The delayed interval is also illustrated in Fig.3A; the applied time shift is indicated with the orange arrow. We computed the lexicon confusion matrix (CM) for any given delay. The confusion matrix indicates which word is reactivated when a target word is presented; the sum of each column is one and corresponds to the fraction of trials (column) assigned to each target (row). Values close to one on the CM’s diagonal indicate correct recognition, while off-diagonal elements indicate mistaken phoneme sequences. The average recognition score is computed from the confusion matrix using Cohen’s *κ* to correct against chance.

#### Average Recognition Delay

The average recognition delay is computed for both the uniqueness and offset measures starting from the *κ*-score. The ARD is the linearly weighted average of the delay function, with respect to the *κ*-scores. That is:

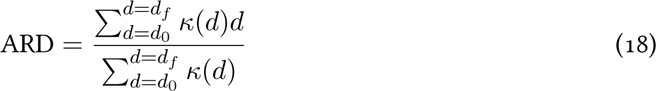

#### Effective connectivity matrix

Excitatory cells have recurrent connections with 20 % of the remaining network’s cells. These connections are formed between the soma and the soma in the point-neuron models and the soma and the dendritic compartment in the dendritic models. We label these connections 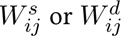, where *s* and *d* are the somatic or dendritic compartment of the post-synaptic neuron *i*, onto which the pre-synaptic neuron *j* projects.

For the asymmetric model 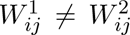 and the two matrices were drawn independently. For the symmetric model 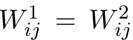, and for the model with a single dendrite only 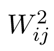 is set to zero. For all the dendritic models, 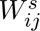 is also set to zero. The opposite is true for the point-neuron models, for which only 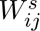 is defined. We define the effective connectivity matrix (C) as the average of the connections between neurons from the pre-synaptic assembly *A* to the post-synaptic assembly *B*

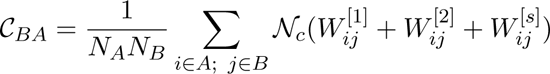

where *N_A_*(*N_B_*) is the number of neurons that belong to the assembly *A* (*B*) and the superscripts in *W* ^1^,^2^*^,s^*, indicates whether the post-synaptic connection targets the two dendrites or the soma. The factor N*_c_* accounts for the compartments that receive inputs, it is 0.5 in the symmetric and asymmetric dendritic models and 1 otherwise. The C is composed of four blocks, the connections between word assemblies (C*^W^ ^→W^*), between phoneme assemblies (C*^P^ ^→P^*), from phonemes to words (C*^P^ ^→W^*), and from words to phonemes (C*^W^ ^→P^*).

#### Block-wise update of the effective matrix

To test which connection types are necessary for word recognition, we leveled the synaptic strength of selected synapses to the initial synaptic strength value (*W*_0_). The synapses are updated based on their pre and post-synaptic cell assemblies. The synaptic update is defined by:

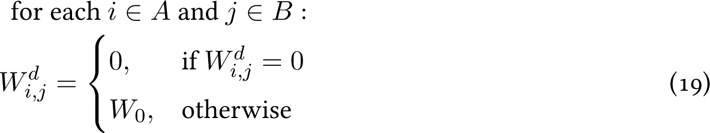

where *i, j* are the indices of two excitatory cells, *A, B* are the post and pre-synaptic assemblies, and the superscript *d* indicates the dendritic compartments, which are updated independently. The update rule in Eq.19 is iterated among all the *A, B* assemblies belonging to the block that has to be flattened. For example, to level the phonemes-to-word connections (C*^P^ ^→W^*), the update rule is repeated for all the assemblies *A* associated with words, and all the assemblies *B* associated with phonemes.

#### Flattening of lexical connections

The strength of the connections between the phonemes and word assemblies depends on the serial position of the phoneme in the word (Results, Fig.6A). To evaluate the computational role of the differences in synaptic weights, we modified 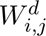 such that all connection strengths from each phoneme to the word populations are the same. The result is an effective matrix in which the connections from any phoneme to the word that contains them have the same strength. To this aim, we used a similar procedure to Eq.19, selecting *A, B* only among the lexical connections. In this case, the synaptic value was not updated with the initial value but with the average of all the synapses targeting the word population 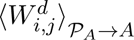 Namely,

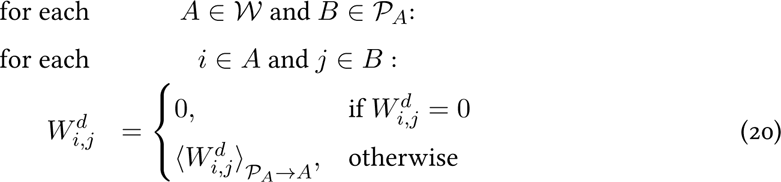

where W is the set of word assemblies and P*_A_* is the set of phonemes contained in the word *A*. The result of the update rule in Eq.20 is illustrated in Fig.6B.

**Table 6:**
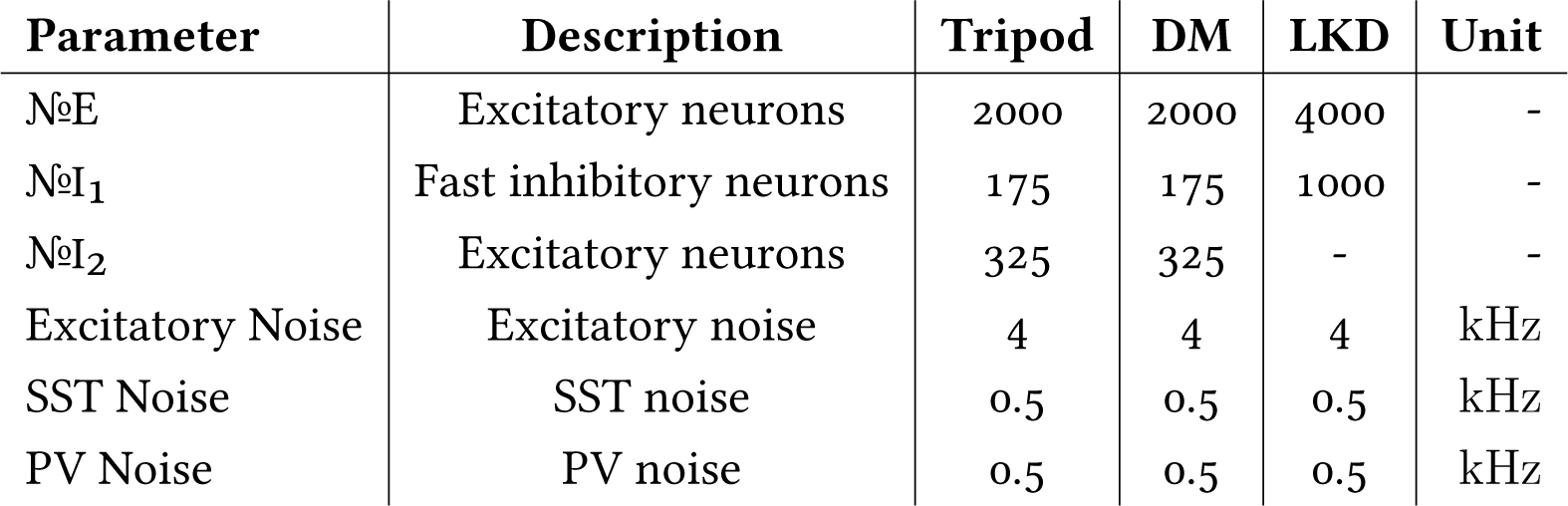
Number and types of neurons for each model, intensity of the background noise.

**Table 7:** Connectivity for the three network models. The connections subject to STDP and iSTDP are in bold text, the synapses are listed in the couple post-pre synaptic and the squared parentheses indicate the post-synaptic target compartment. The numbers indicate respectively the connection probability and initial synaptic strength. The synaptic strength is in parenthesis, expressed in pF.

| Synapse | LitwinKumar & Doiron | Duarte & Morrison | Tripod |
| --- | --- | --- | --- |
| $E[d]E$ | - | - | 0.2 (10.78) |
| $E[s]E$ | 0.2 (2.76) | 0.168 (0.45) | - |
| $I_1E$ | 0.2 (1.27) | 0.575 (1.65) | 0.2 (5.27) |
| $I_2E$ | 0.2 | 0.244 (0.638) | 0.2 (5.27) |
| $E[s]I_1$ | 0.2 (48.7) | 0.6 (5.148) | 0.2 (15.8) |
| $E[d]I_2$ | - | - | 0.2 (15.8) |
| $E^sI_2$ | - | 0.465 (4.85) | - |
| $I_2I_2$ | - | 0.381 (0.83) | 0.2 (16.2) |
| $I_1I_1$ | 0.2 (16.2) | 0.55 (2.22) | 0.2 (16.2) |
| $I_1I_2$ | - | 0.379 (1.47) | 0.2 (1.47) |
| $I_2I_1$ | - | 0.24 (1.4) | 0.2 (0.83) |

**Table 8:** Lexica that are used in the simulations. The table presents the words that compose each lexicon, the average phonological overlap, and the lexicon size.

| Vocabulary | Words | Phon. Overlap | Lexicon size |
| --- | --- | --- | --- |
| Identity | a, b, c, d, e, f, g, h, i, j | 1 | 10 |
| No overlap | abcd, efghi, jkl, mno, pqrst, uvw, xyz | 1 | 7 |
| TIMIT | all, ask, carry, dark, don't, had, like, me, oily, rag, she, suit, that, wash, water, year, your | 2.63 | 17 |
| Digits | zero, one, two, three, four, five, six, seven, eight, nine | 3.33 | 10 |
| Overlap | pollen, gold, golden, doll, lop, god, log, poll, goal, dog | 5.0 | 10 |
| Cohort | capias, capillary, capistrate, capital, capitalise, capitalism, capitalist, capitate, capitol, capitoline, capitulate, capitulation | 7.69 | 12 |
| TISK | art, artist, arts, rats, star, start, stat, tar, tars, trist | 8.33 | 10 |

## Appendix A Network activity during the associative and recall phase

To characterize network activity, we measured the average firing rate and the variability in the interspike intervals (ISI)(Fig.2*B*). We observe the firing rate of the excitatory neurons is heterogeneous and ranges from 0 Hz to 15 Hz. The rate of the excitatory population (Tripod neurons) divides into two groups. One group comprises neurons that receive external stimuli and thus belong to an assembly (*Exc. with inputs*); these cells fire at various rates in the range 5 Hz to 15 Hz. The second class is composed of neurons that do not receive external stimuli; they fire at a lower rate, in the range 0 Hz to 10 Hz. Because of the heterogeneity in the neurons’ rate, we cannot use the standard coefficient of variation for the ISI measure, it would associate higher variability to neurons with higher rates (Holt *et al*., 1996). For an unbiased estimate of the ISI CV we computed the coefficient of variation based only on adjacent interspike intervals (ISI CV2). Accordingly, the differences between excitatory neurons are less visible in the ISI variability; neurons with no inputs have ISI CV2 in the interval 0.5 to 1.3 while the measure is slightly lower for neurons that receive external stimuli. Finally, interneurons of types I1and I2also fire at heterogeneous rates in the interval 5 Hz to 50 Hz, their coefficient of variation remains around 1, consistent with Poisson input they receive from the remainder of the network.

The variability observed in the rate of excitatory neurons is unexpected. Indeed, the rapid compensatory mechanism implemented in the network (iSTDP on somatic compartments) should steer the neurons to fire at 10 Hz. In point-neuron models (both 4 and Litwin-Kumar & Doiron (2014)), the rate is tightly centered at the homeostatic value, although some neurons receive inputs, and some do not. These initial results show that including dendrites in the neuron model greatly impacts the network dynamics and leads to heterogeneity in the firing rate in the face of counteracting homeostatic mechanisms.

We exclude the presence of pathological synchronicity, we computed the fast Fourier transform (FFT) of the network firing rate during the associative and recall phases. At this scope, the spikes of the excitatory and inhibitory populations were merged and binned in intervals of 5 ms, corresponding to Nyquist frequency of 100 Hz for the FFT. The spectrogram for the recall phase is illustrated in App. Fig.1 *C*. The frequency analysis reveals clear peaks at the frequency of the phonemic input 20 Hz and 40 Hz - corresponding to the fundamental frequency and the first harmonic of the phonemic stimuli (50 ms). The peaked spectral reveals the resistor-capacitor (RC) circuit subserving the Tripod neuron’s dendritic compartments; the dendritic compartments start charging when the phoneme projections activate and slowly decay when it is turned off. A smaller but more interesting peak is visible around 5 Hz (orange band in the panel); this frequency interval corresponds to word inputs. Because no external activity is carried out on the word projections in the recall phase, the peak in the spectrum must correspond to the reactivation of the word assemblies, indicating an internally generated response. The fact that words are interleaved by 50 ms of silence (absence of inputs) offers a possible confound, however, the peak at low frequency is present also when there is no silence among words in the recall phase.

Additional evidence that the word assemblies populations are reactivated in the recall phase is obtained by the comparative analysis of the firing rate correlations. In App. Fig.1 *D*, we illustrate the average autocorrelation function of network subpopulations corresponding to phonemes, words, or randomly sampled neurons (control group of 100 neurons); we mark the half-height (HH) correlation time, that is, the time for the firing rate to decorrelate of 50 %. For the phonemes populations, the differences between the associative and recall phases are expected to be minor because they receive the same external input in the two phases; the HH is 48 ms in the associative phase and (55 ms) in the recall. Despite being a marginal difference, the HH in the recall phase is systematically (10 network samples) longer than in the associative phase; we hypothesize this to be due to the fewer external inputs in the former condition, which interfere less with the reverberations of the assemblies. The word assemblies shows a radically different scenario. In the early phase, the populations are maintained in continuous firing by the external projections, lasting 150 ms to 350 ms. The assemblies have HH correlations longer than 100 ms (orange dashed lines). When the external stimulation is removed (recall phase, orange solid lines), the correlation time of the word assemblies shrinks to 75 ms. The difference in the two conditions indicates that the retrieved activity is less stable than the input activity. Nonetheless, the HH of word populations is twice the HH of the control populations (grey, 40 ms), which points to the role of the acquired word engram in fostering activity in the assemblies.

**Appendix Figure 1:**
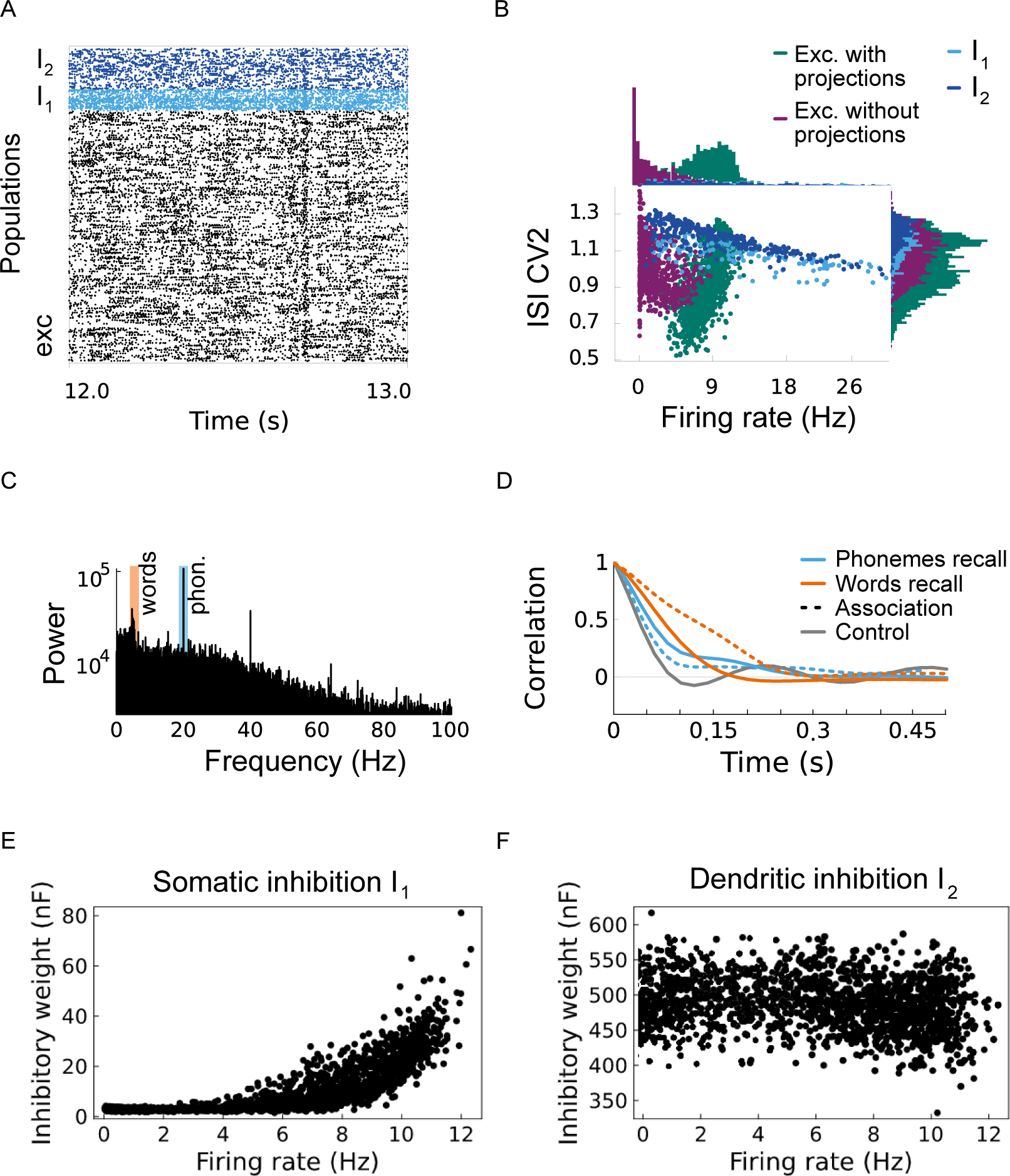
Firing activity during the associative and recall phase. (**A**) Raster plot of the Tripod network during an interval of 1 s. Excitatory dendritic neurons are shown in black, fast-spiking interneurons (I1) in light blue, and slow-spiking interneurons (I2) in dark blue. (**B**) Mean firing rate and CV2 ISI for excitatory and inhibitory populations. Excitatory neurons are divided into two groups, those with (green) and without (purple) external input. The former have a higher firing rate and a lower CV2 ISI. (**C**) The frequency spectrum of the excitatory firing activity across the entire simulation (5 min). The large peak at 20 Hz and 40 Hz reflect the frequency of the phonemic inputs (blue marker). Conversely, the smoother peak at low frequency (orange marker) is due to the network’s internal dynamics. The spectrogram is computed for a recall protocol with no silence in between each word, hence the peak is due to word re-activation, Overall the frequency spectrum indicates the network is in the asynchronous firing regime. (**D**) Assembly firing rate autocorrelation averaged over phonemes (blue) and word (orange) assemblies in the associative (dashed) and recall phase (solid). The grey line illustrates the autocorrelation of a population of randomly sampled neurons. The word assemblies (orange) have the longest autocorrelation time in both phases. It indicates that the engram acquired during the association phase can collectively reactivate and sustain firing during retrieval.

Finally, we report the strength of the inhibitory neurons with respect to the firing activity of the excitatory cells. The difference in the inhibitory plasticity rule decouples the two inhibitory populations. In the case of the I1, the synaptic strength correlates with the firing rate of the Tripod neurons. Inhibitory synapses are small for cells that seldomly fire, and strong for those that fire often App. Fig.1 *E*. Conversely, the dendritic inhibition does not depend on the rate, all excitatory cells form strong inhibitory synapses on the dendritic compartments App. Fig.1 *F*.

## Appendix B Formation and maintenance of word memories

**Appendix Figure 2:**
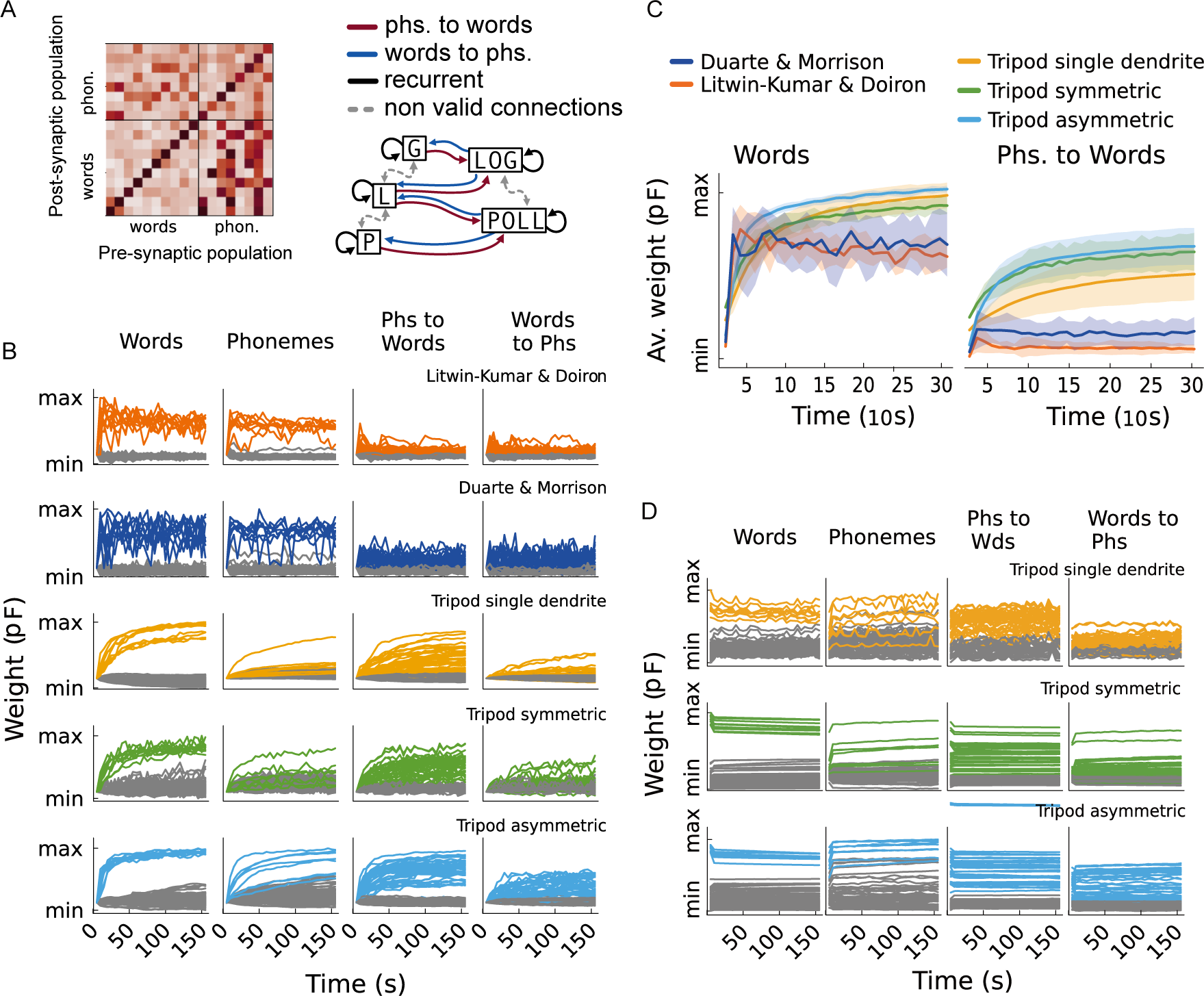
Formation and maintenance of network memories. (**A**) Effective connectivity matrix for the Tripod Asymmetric model, the scheme illustrates the types of connections analyzed in the remainder of the figure. (**B**) Weights formation during the associative phase, for all the connections in *A* and the five analyzed models. (**C**) Average recurrent and feedforward connection weights during the memory formation phase. The five models are drawn together for comparison. The maximum and minimum refer to the extrema of the EC for each model. (**D**) Weights decay during the recall phase for the three dendritic models. The synaptic strengths remain overall stable with the largest drop happening in the word recurrent connections.

We can visualize the evolution of the recurrent and forward connections by computing the effective connectivity matrix throughout the simulation (C, App. Fig.2 *A*). The panels in App. Fig.2 *B* show the average synaptic strength for each element of the effective connectivity matrix during the associative phase. Each row indicates one of the five models tested on the *Overlap* lexicon. Similarly to the main text, the maximum and minimum values refer to the maximum of the effective connectivity matrix and the initial synaptic weight.

All models have strong recurrent word memories, only the dendritic models have distinct phonemes-to-words and word-to-phonemes connections. In addition, the strengthening of synaptic connections on the dendritic compartments is slower than in the soma models. Synapses grow smoothly throughout the associative interval. Conversely, in the point neuron models, the recurrent memories are more volatile, and prone to be overridden, as indicated by the large fluctuations in the average synaptic weights. A more compact view of the memory formation process is presented in App. Fig.2 *C*, with a direct comparison of the recurrent and feedforward connections of the five models. For all the dendritic models, the learning trajectory is exponentially fast, with a timescale in the order of 10 s for the recurrent memories and 100 s for the feedforward connections. In contrast, the two point-neuron models reach high synaptic strength for the recurrent connections within just a few words presentation, but the non-recurrent connections remain close to the initial synaptic weight.

To verify the stability of the memory formed in the associative phase, we run the network with excitatory plasticity active during the recall phase. The panels in App. Fig.2 *D* show the evolution of the synaptic strengths over 150 ms during the presentation of the phoneme sequences. The weights do change less than 5 % from those reached at the end of the associative phase; the recognition score, not shown, also remained high, despite a systematic drop of a few percent points. In the main text, the weights were instead frozen because the integration of the STDP equations is computationally demanding for the simulation. We have not tested whether the weights would be maintained in the absence of stimuli, from the results in Appendix A, however, we estimate that the synchronized burst associated with the network could interact negatively. However, the presence of structured connections, learned in the associative phase, could also favor the reactivation of words and preserve the weights. Future work should take a closer look at the maintenance of the connectivity against the presentation of new stimuli or silence.

## Supplementary Material

**SI Figure 1 SI Figure 2:**
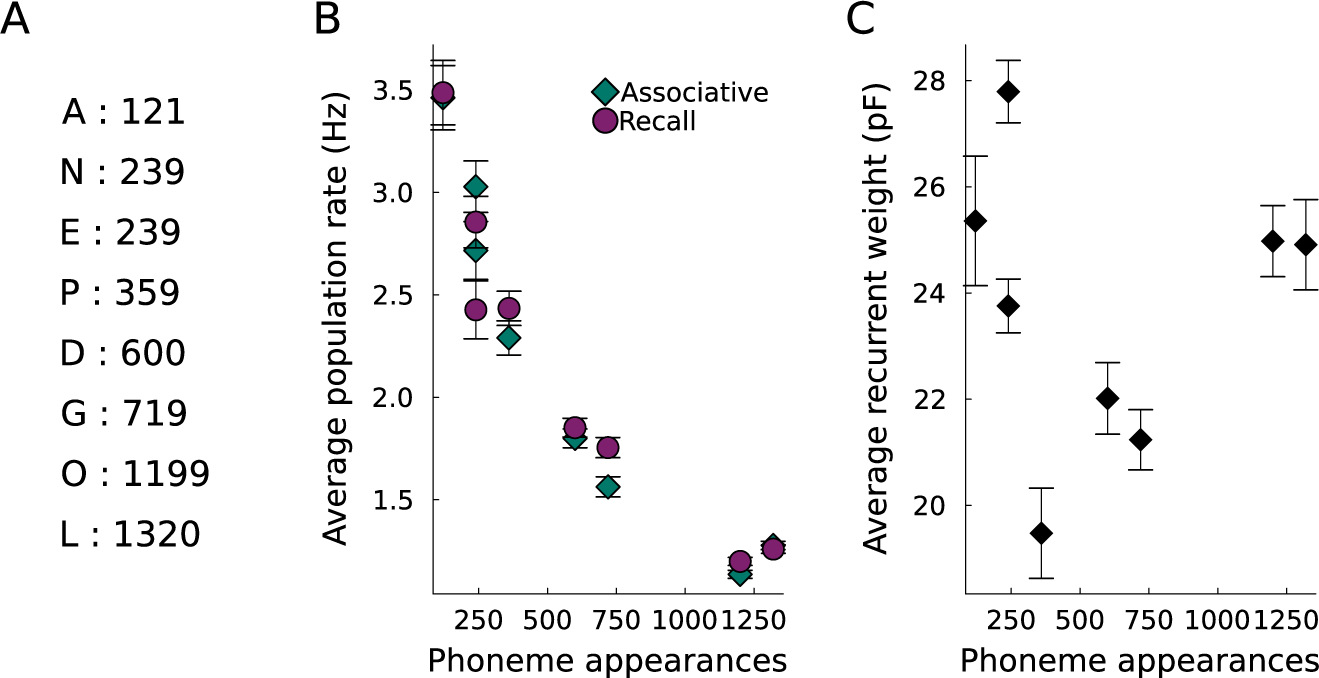
Activity and recurrent weights of phoneme assemblies in the Overlap lexicon. (**A**) Frequency of the phonemes in the input sequence. (**B**) More frequent phonemes have low firing rates, possibly due to neuronal adaptation of the firing rate. (**C**) Additionally, the synaptic scaling applied to glutamatergic synapses tends to reduce the synaptic strength of phonemes that are contained in several words. Because the incoming synaptic weight is fixed, phoneme populations that occur in multiple words must share their synaptic strength among more pre-synaptic pathways, reducing the available synaptic resources for the recurrent connections.

**SI Figure 3:**
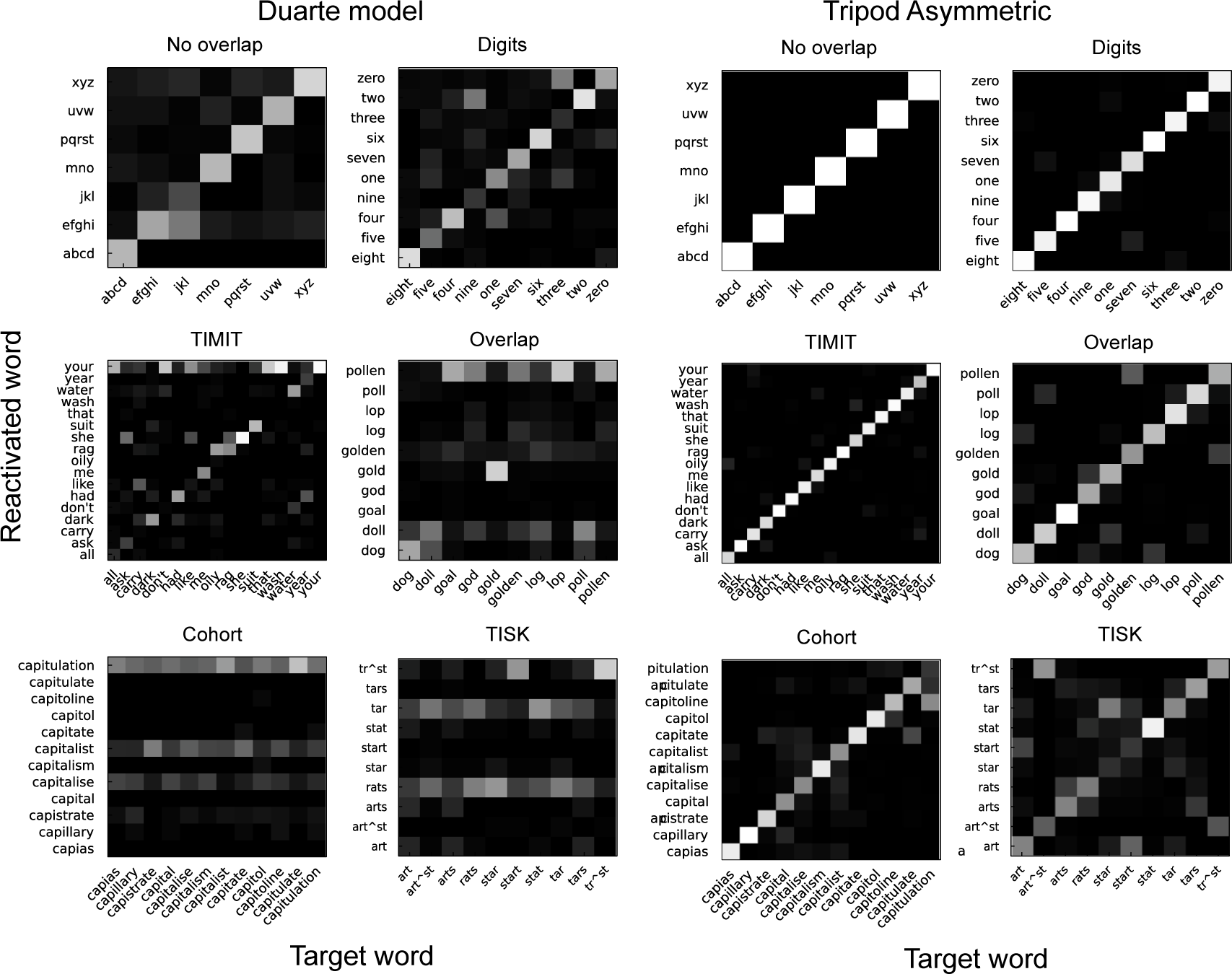
Confusion matrices for point neuron and dendritic models. Confusion matrices for six of the lexica that were measured. The left columns show the confusion matrix for the Duarte model, and the right columns those for the asymmetric tripod network. The elements on the diagonal indicate correct word recognition; conversely, the off-diagonal ones indicate wrong recollection. E.g., in the dendritic network, when tested in the Cohort lexicon, the word *capitulation* is often confused for the word *capitoline*, but the opposite does not happen. Crucially, the confusion matrices of the point neuron model show correct word recognition for the lexicon with less phonological overlap but are completely random for the three with large phonological overlap. This is not the case for the dendritic network; despite the matrices are

**SI Figure 4:**
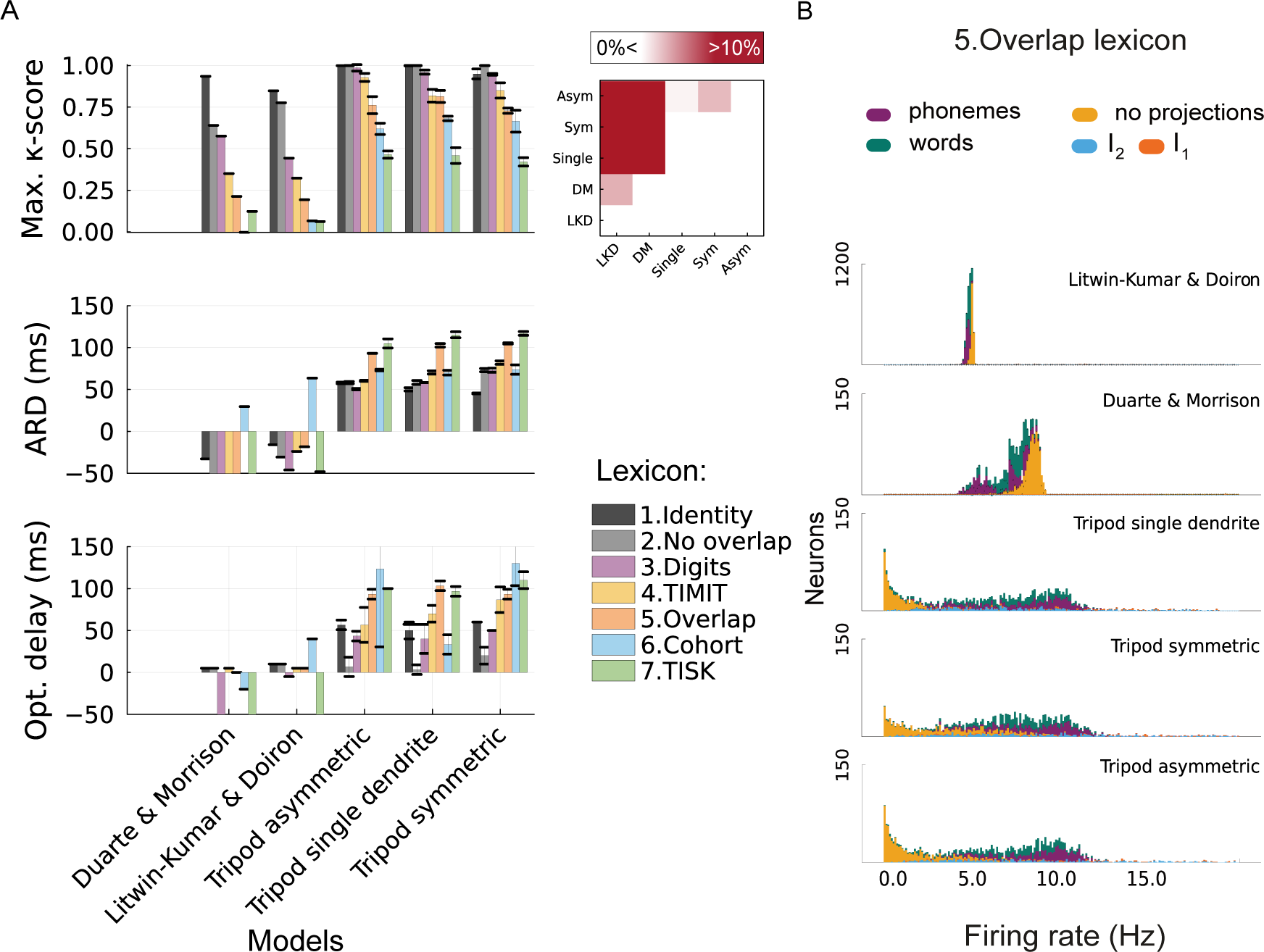
Model comparison for the full interval measure firing profiles. (**A**) Comparison of recognition scores and average delay for the full interval measure. The panel up-right shows the average score differences between the five models; asymmetrical and symmetrical models perform slightly better than the single dendrite model. (**B**) Firing rate profile for the five network models during the recall phase of the Lexicon dictionary. The point neuron models have distributions peaked around the firing rate promoted by the inhibitory plasticity; there is a minor difference between the neurons within assemblies and those not. In contrast, the firing rate of dendritic models is largely heterogeneous, and neurons with no direct projections have a low firing rate. Remarkably, the firing rate profile is similar for the three dendritic models. We hypothesize that heterogeneity in the average firing rates of the neurons is an indicator of the network performing the task correctly.

**SI Figure 5:**
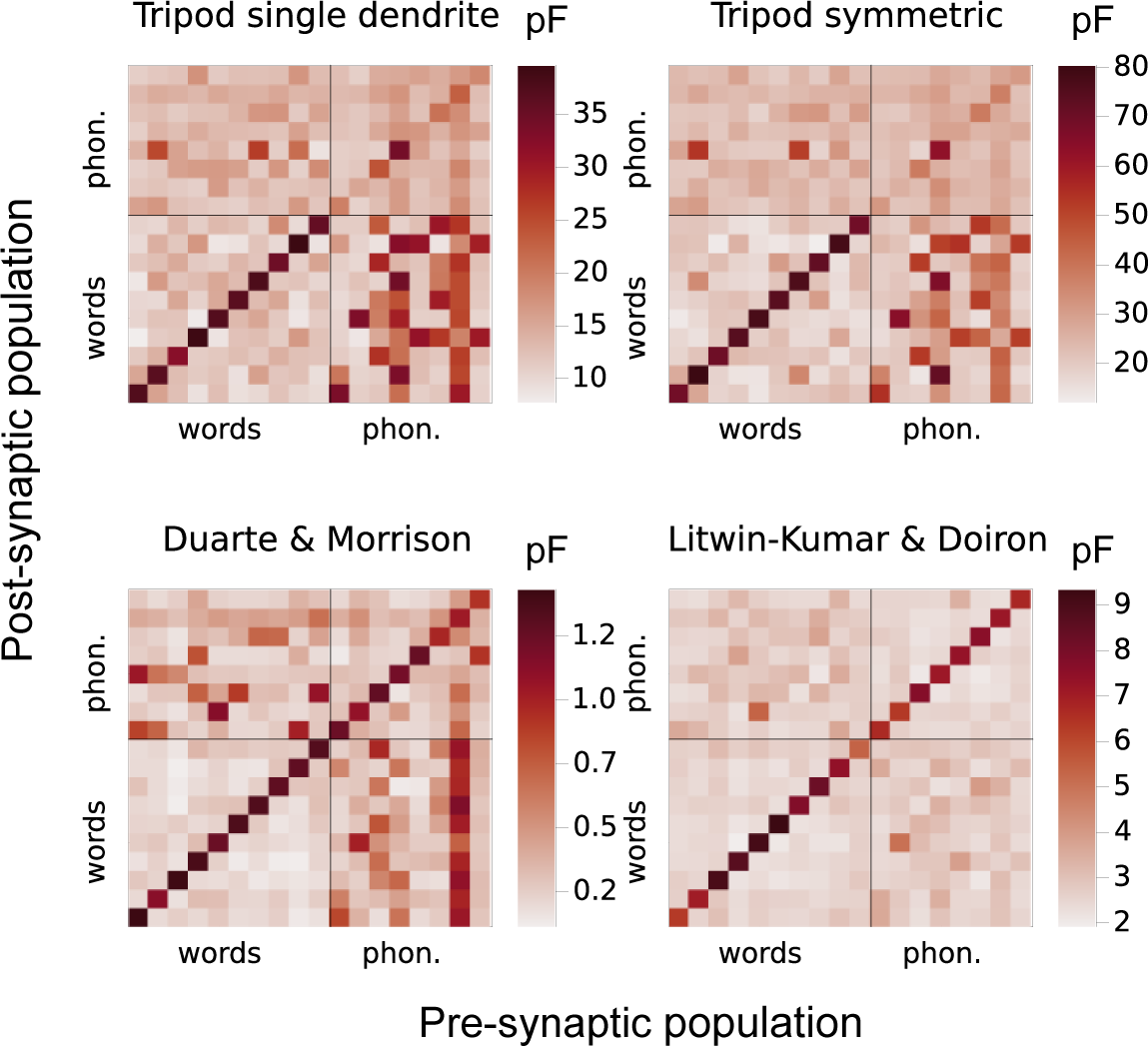
Effective connectivity matrices for each model in the *Overlap* lexicon. Effective connectivity matrices for the symmetric and single dendritic models (the asymmetric model is portrayed in Fig.5)*A* and the two-point neurons models (bottom). The dendritic models develop strong connections between phonemes and word assemblies (bottom-right quadrants). In contrast, the point neuron models do not - as in the LKD network-develop connections that are ineffective in re-activating the word populations - as in the DM.

**SI Figure 6:**
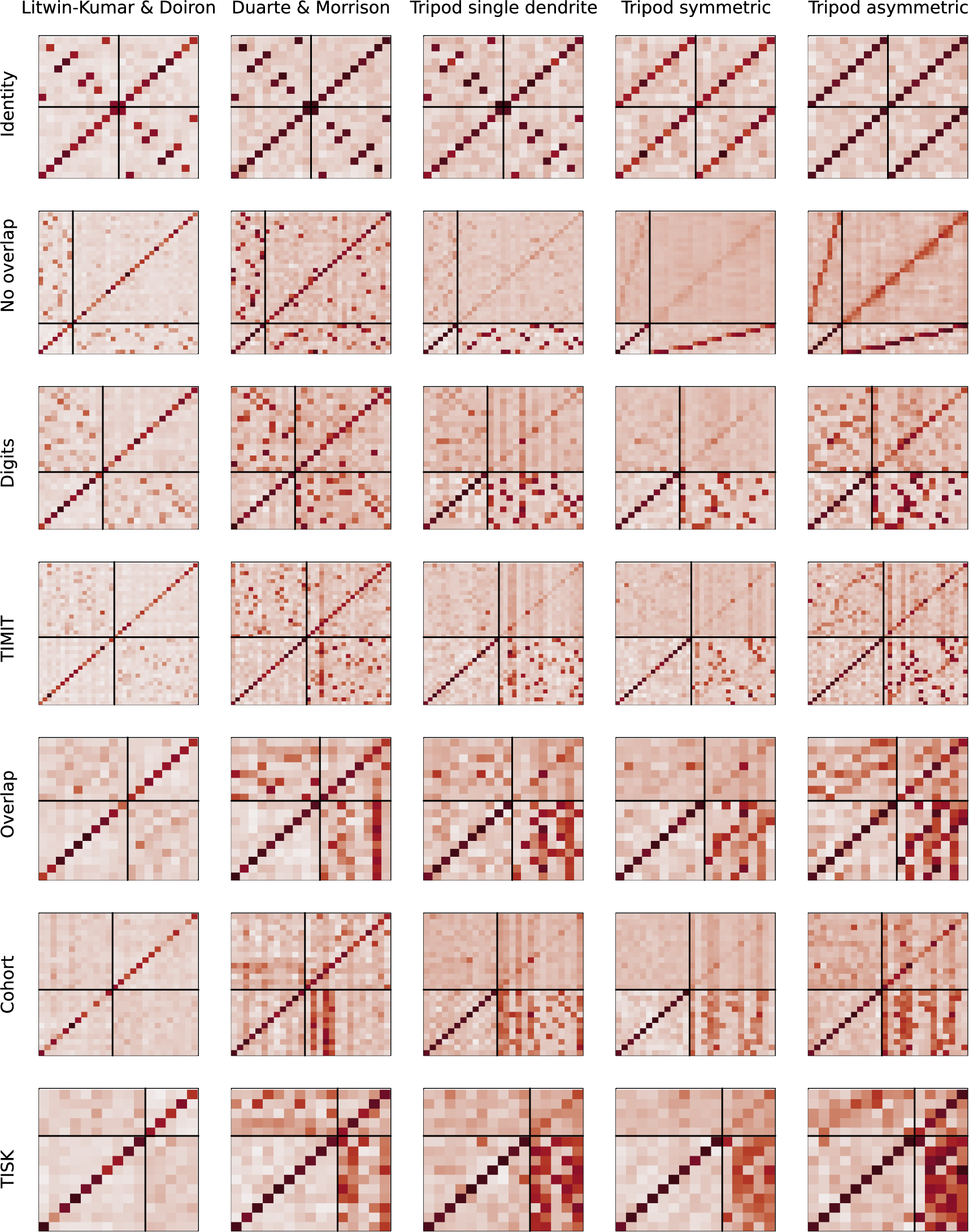
Effective connectivity matrices for all models and lexica. The overall comparison of the effective connectivity matrices indicates that point-neuron models can form hetero-associative connections between phonemes and words (bottom-left quadrants) when the recollected words have no phonological overlap (first two rows, *Identity* and *No overlap* lexica)

**SI Figure 7:**
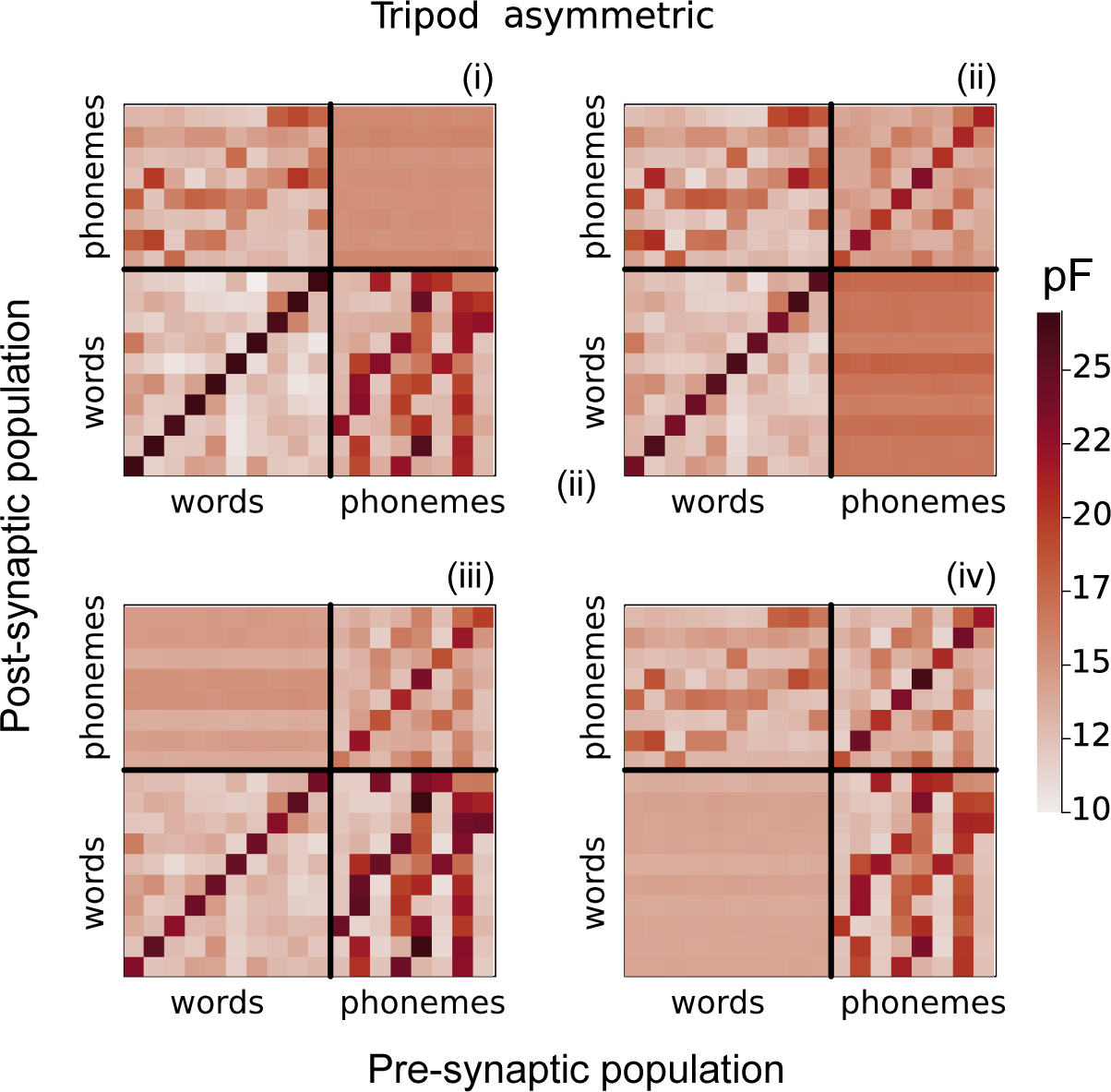
Confusion matrices for point neuron and dendritic models. Confusion matrices for six of the lexica that were measured. The left columns show the confusion matrix for the Duarte model, and the right columns those for the asymmetric tripod network. The elements on the diagonal indicate correct word recognition; conversely, the off-diagonal ones indicate wrong recollection. E.g., in the dendritic network, when tested in the Cohort lexicon, the word *capitulation* is often confused for the word *capitoline*, but the opposite does not happen. Crucially, the confusion matrices of the point neuron model show correct word recognition for the lexicon with less phonological overlap but are completely random for the three with large phonological overlap. This is not the case for the dendritic network; despite the matrices are

**SI Figure 8:**
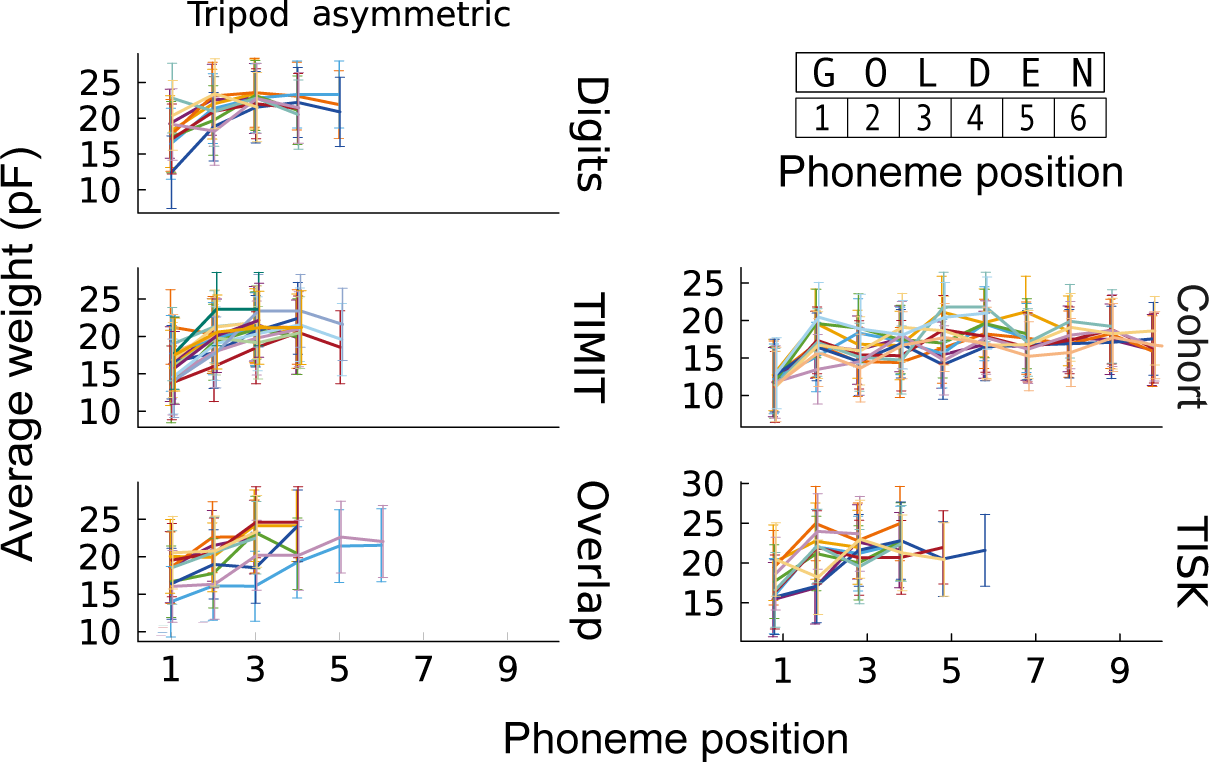
Synaptic efficacy of phoneme to word connections. The synaptic strength between phonemes and word populations is compared with the serial position of the phoneme in the word. The five panels show the phonemes-to-word connectivity for all the lexica that require memory, for the Tripod asymmetric model. The measures are averaged together in Fig. 5 of the main text. Conversely, here, the scale changes for every model.

**SI Figure 9:**
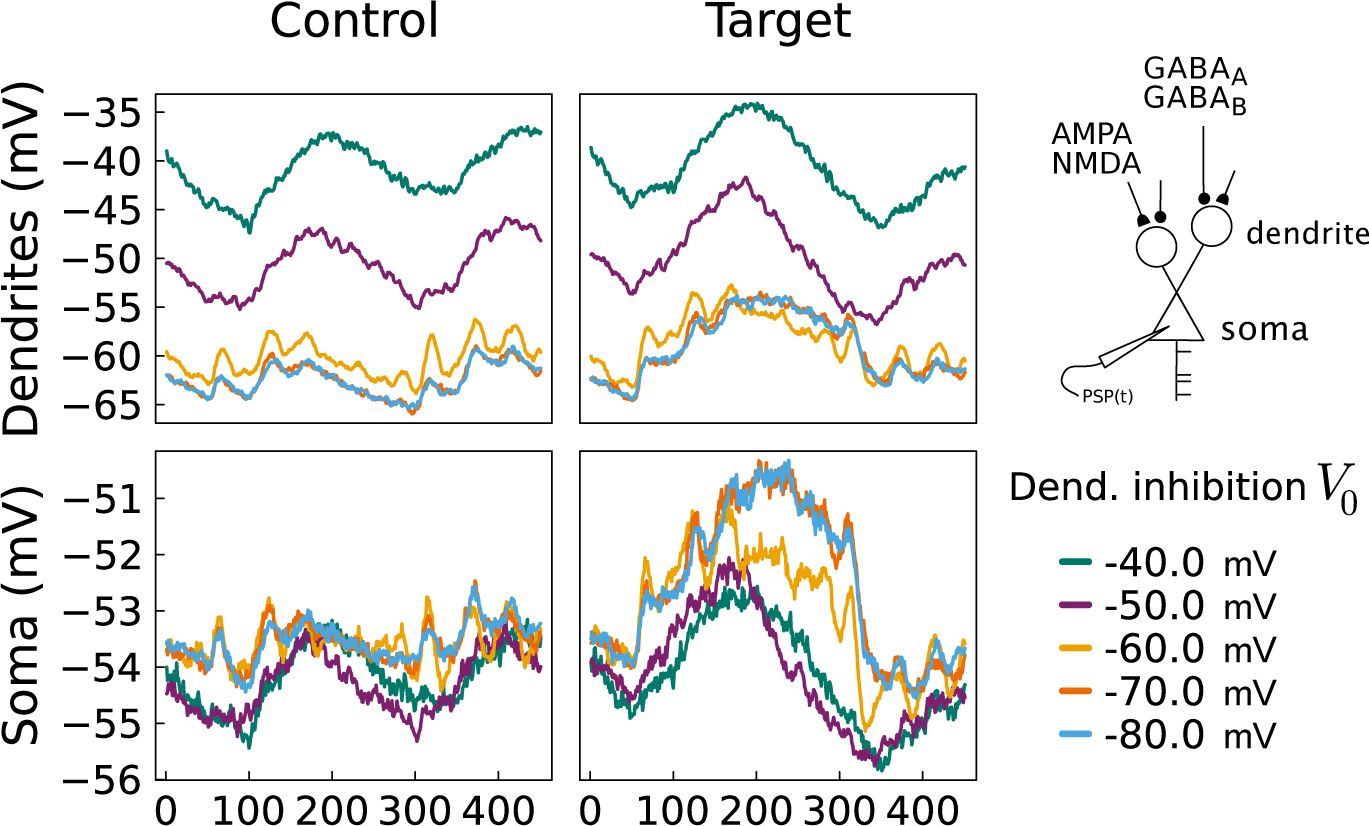
Dendritic inhibition governs signal-to-noise ratio in dendritic membrane potential. (**A**) The average membrane potential of the dendritic and somatic compartments of a word assembly during the presentation of a phoneme sequence. The left panel shows the average potential of 100 cells randomly selected cells (control), instead, the right panel illustrates the membrane potential of the word assembly corresponding to the phoneme sequence presented (target). The colors indicate five different target membrane potentials (*V*_0_) used for the v-iSTDP rule. The plot shows that for more hyperpolarized target potential (−70 mV to −80 mV) the difference between target and control increases.

